# Strong Selection, but low repeatability: Temperature-specific effects on genomic predictions of adaptation

**DOI:** 10.1101/2024.09.12.612579

**Authors:** Alexandre Rêgo, Julian Baur, Camille Girard-Tercieux, Maria de la Paz Celorio-Mancera, Rike Stelkens, David Berger

## Abstract

Climate warming is threatening biodiversity by increasing temperatures beyond the optima of many ectotherms. Due to the inherent non-linear relationship between temperature and the rate of cellular processes, such shifts towards hot temperature are predicted to impose stronger selection compared to corresponding shifts toward cold temperature. This suggests that when adaptation to warming occurs, it should be relatively rapid and predictable. Here, we tested this hypothesis from the level of single-nucleotide polymorphisms to life-history traits, by conducting an evolve-and-resequence experiment on three genetic backgrounds of the beetle, *Callosobruchus maculatus*. Indeed, phenotypic evolution was faster and more repeatable at hot, relative to cold, temperature. However, at the genomic level, adaptation to heat was less repeatable when compared across genetic backgrounds. As a result, genomic predictions of phenotypic adaptation in populations exposed to hot temperature were accurate within, but not between, backgrounds. These results seem best explained by genetic redundancy and an increased importance of epistasis during adaptation to heat, and imply that the same mechanisms that exert strong selection and increase repeatability of phenotypic evolution at hot temperature, reduce repeatability at the genomic level. Thus, predictions of adaptation in key phenotypes from genomic data may become increasingly difficult as climates warm.

## Introduction

Whether evolution is repeatable, and whether the same genes contribute to recurrent phenotypic adaptations, is a long-standing question with fundamental implications for predicting evolutionary responses to changing environments [1–3]. Recent studies have shown that, while evolution at the phenotypic level often is repeatable, the genes that contribute to adaptation can be highly contingent on the evolutionary history of populations [2, 4, 5]. Nevertheless, theory [6] and recent evolve and re-sequence (E&R) experiments [2, 7–13] have revealed that broad-scale patterns also emerge at the genomic level; adaptation is often polygenic, and while individual DNA polymorphisms are unlikely to repeatedly contribute to adaptation, the same molecular pathways are often involved, sometimes even over long evolutionary timescales [14]. This type of information is necessary to gain an informed opinion about adaptive potential under climate change [1, 15, 16], and is becoming increasingly important given the growing reliance on genomic data to inform conservation practices [17–20]. However, whether genomic estimates alone can accurately predict extinction risk and adaptation in key ecological phenotypes remains poorly understood [3, 21, 22].

Several factors can influence the probability of observing repeated evolutionary responses, such as differences in standing genetic variation in natural populations [23–25], or previous differentiation between evolving lineages that may impact the fates of segregating alleles whose fitness effects depend on allele frequencies at other loci (i.e., epistasis) [26–28]. Theory additionally suggests that environmental changes that impose stronger selection are more likely to produce repeatable evolutionary outcomes compared to environments that impose weak selection [2, 6]. However, despite being critical for eco-evolutionary forecasting, the relative importance of these factors in producing (non)repeatable outcomes of evolution are not well understood [3, 29, 30].

Genetic adaptation to temperature is becoming evermore crucial under contemporary climate change. To what extent high and low temperatures result in repeatable evolutionary outcomes, and whether changes at the genomic level are informative of phenotypic adaptation, can further inform the future management of threatened populations. However, studies which incorporate both phenotypic and genomic information on thermal adaptation are rare [31, 32]. Yet, cases of adaptation to temperature are amenable to exploring the causes of evolutionary repeatability due to a fairly deep understanding of thermal physiology [33, 34]. For example, thermodynamic constraints on cellular processes often manifest as asymmetries in thermal performance curves of ectotherms, where performance measures more rapidly decline at hotter than colder temperatures at equal distances from a thermal optimum [35] (Fig. 1A). Organisms far-displaced from their thermal optimum towards hotter temperatures may thus experience strong increases in selection for beneficial alleles, suggesting that evolution may generally be faster and more predictable in warming climates [36–38]. Understanding the evolutionary consequences of this physiological temperature-dependence is thus of central importance for evaluating the prospects of employing genomics data to predict phenotypic adaptation and extinction risk under future climate change.

**Figure 1:**
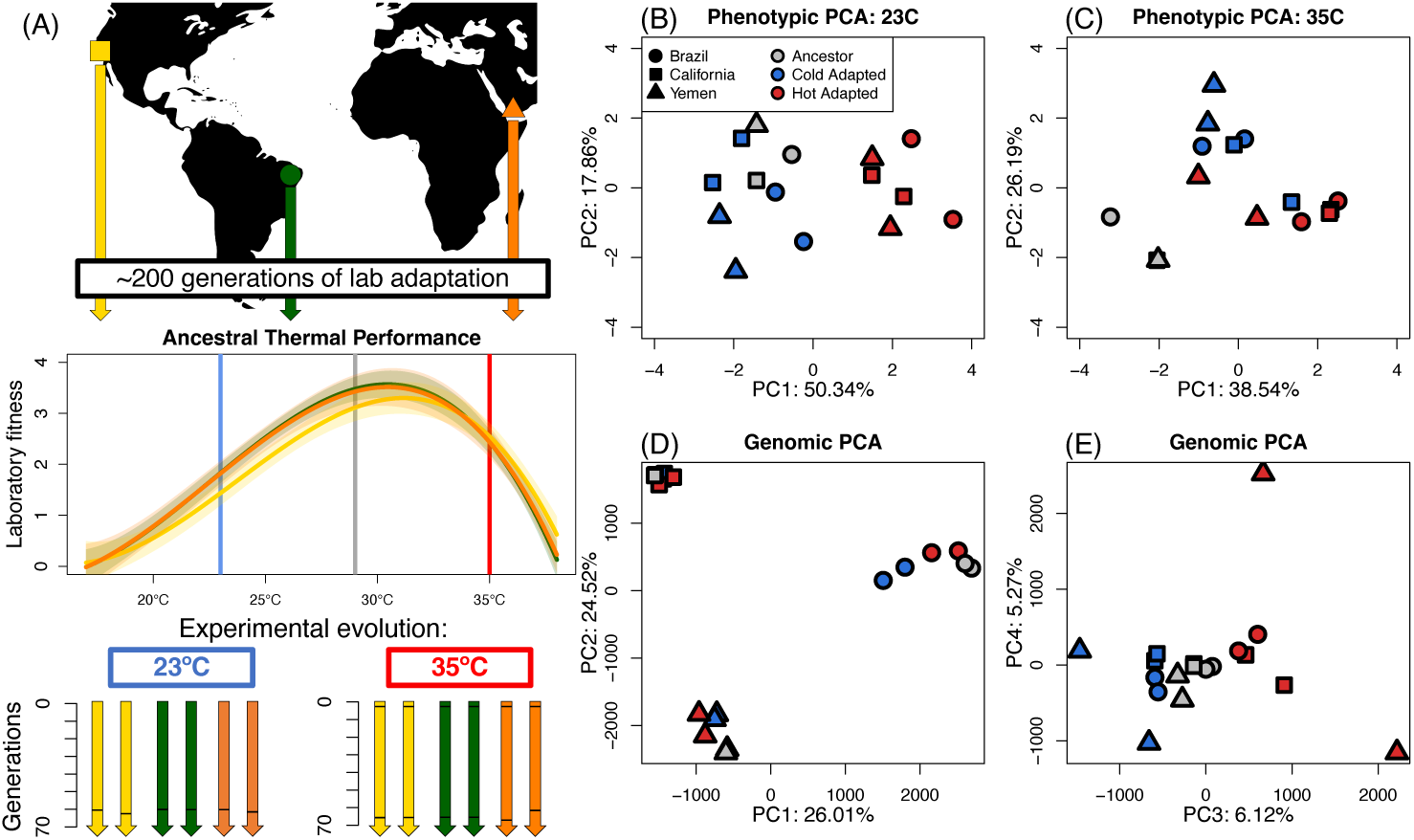
Experimental design. (A) Populations were collected from California (yellow), Brazil (green), and Yemen (orange) and adapted to benign lab conditions (29°C) for approximately 200 generations. Thermal performance curves of the three populations show the classic asymmetric shape and depict how population growth rate (adult offspring production/generation time) changes with temperature. While mean growth rates are slightly lower at 23°C compared to 35°C, the slope of the performance curve is steeper at temperatures higher than the optimum. To start experimental evolution, lines were placed at either 23°C or 35°C, with two replicates in each treatment per genetic background. Horizontal black bars in the arrows depicting each experimental evolution line indicate the time at which lines were sequenced. The sequencing of heat-adapted lines at generation 3 represent the genomic estimates of ancestral variation. For both the seven life-history phenotypes (B-C) and for all genomic SNPs (D-E), PCA summaries are presented of evolutionary change between ancestral (grey) and heat (red) and cold (blue) adapted lines. Phenotypes were measured in a common garden design including both the low and high temperature.

Here, we combine data on phenotypic and genomic change from replicate experimental evolution lines of the seed beetle *Callosobruchus maculatus*, deriving from an evolve-and-resequence experiment designed to quantify the role of selective determinism versus historical contingency in thermal adaptation. Replicate lines, established from three far-separated geographic populations, were placed at cold (23°C) or hot (35°C) temperature equidistant from their ancestral temperature (29°C) (Fig. 1A). To test for evolutionary repeatability, we estimated evolutionary trajectories at the phenotypic level among seven life-history traits, and at the genomic level across putatively selected SNPs and genes. We then compared evolutionary responses between line replicates (within genetic backgrounds) and geographic populations (between genetic backgrounds) to understand the role of historical contingency and temperature-specific selection on the repeatability of evolution. Finally, we estimated the correspondence between genomic and phenotypic estimates to evaluate the power of genomic data to predict the level of (mal)adaptation.

## Results

### Phenotypic Evolution is faster and more parallel at hot temperature

We first re-analyzed data on thermal adaptation in seven life-history traits of females [39], including three core traits: lifetime offspring production, adult weight and juvenile development time (Fig. 2A), as well as four rate-dependent traits measured over the first 24 hours of female reproductive lifespan: weight loss, water loss, early fecundity and mass-specific metabolic rate (Fig. 1B-C, Fig. S1A). By comparing each of the 12 evolution lines to their respective ancestor, we assessed the direction and magnitude of phenotypic evolution at 23°C (comparing ancestors to cold lines) and 35°C (comparing ancestors to hot lines). We quantified the repeatability of this evolution at each temperature as geometric angles between evolutionary change vectors in multivariate trait space, where an angle of 90 degrees signifies uncorrelated responses and angles of 0 and 180 degrees correspond to perfectly parallel and anti-parallel responses, respectively. Comparisons were made between pairs of evolution lines deriving either from the same or different genetic backgrounds. We also quantified whether populations were converging or diverging over the course of the experiment by comparing the distance between populations at the start (*S_d_*) of experimental evolution (i.e. between ancestors) to the distance at the end (*E_d_*) (i.e. between the evolved lines).

**Figure 2:**
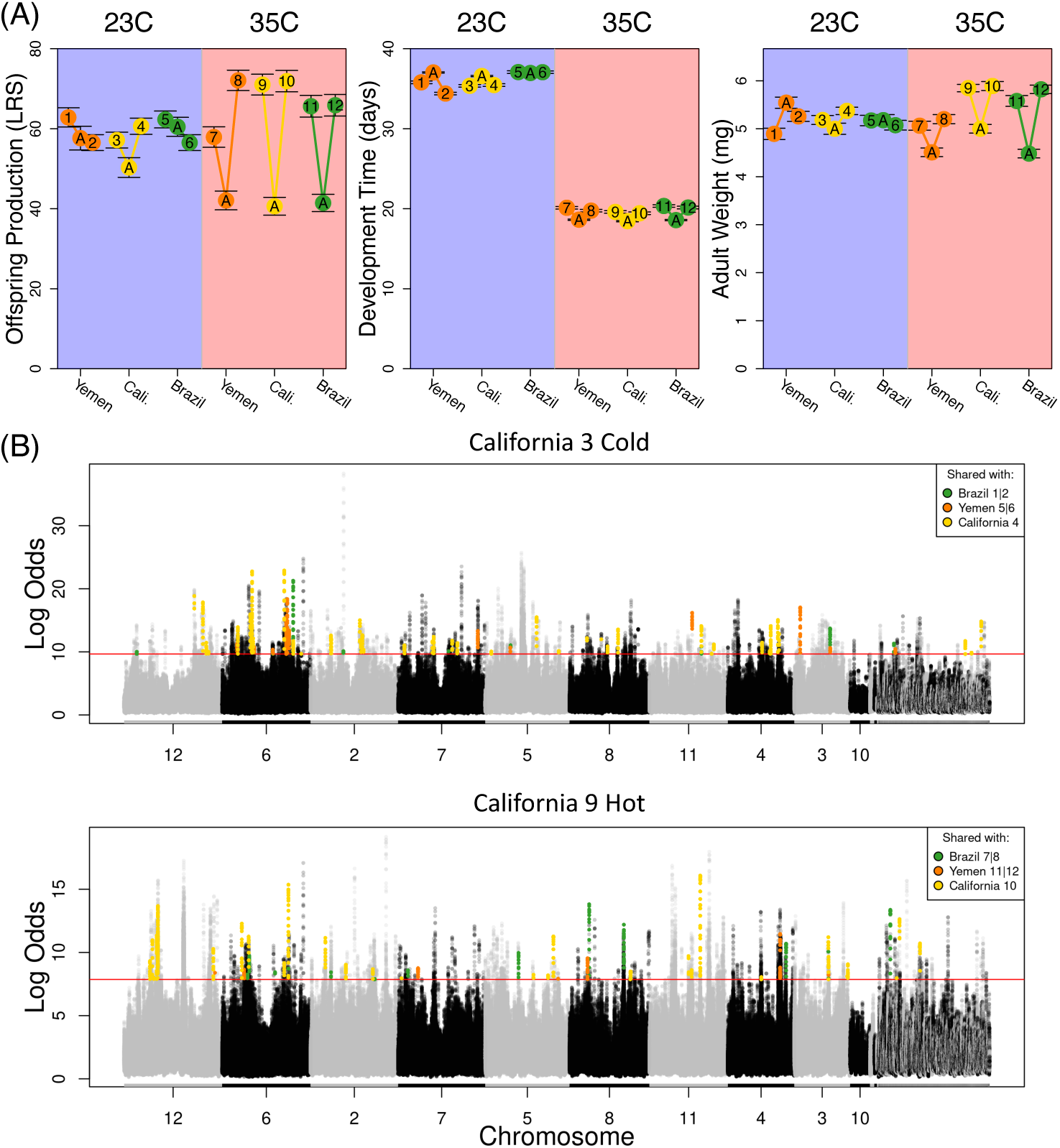
Phenotypic and genomic data. (A) Means (and one standard error) are shown for ancestors and evolved lines for 3 of the 7 measured traits (see Fig. S1 for all traits). Phenotypes of evolved lines (1–12) are shown only as measured in their local temperature regime (cold lines, 1–6, measured at 23°C; hot lines, 7– 12, measured at 35°C). (B) Manhattan plots of rolling average log odds p-values (window size = 20 SNPs) for one heat-adapted and one cold-adapted line from the California background (see Fig. S2 for all populations) across all putatively selected sites (n=475,194) using R package RcppRoll (v.0.3.0). The largest 10 scaffolds (i.e., chromosomes) are labelled. The 0.001th quantile of rolling average p-values is given by the horizontal red line for each line. SNPs whose rolling average p-values are also above their respective 0.001th quantile in other populations are colored. There is a clear polygenic signal of adaptation. Note that the California lines shown here exhibit many more SNPs in common with the single other California replicate (yellow points) than those shared with any of the four line replicates of different origin (green and orange points).

Hot lines exhibit higher evolutionary rates per-generation (∥*x̄*_35_∥ = 0.87 ± 0.14) compared to cold lines (∥*x̄*_23_∥ = 0.5 ± 0.07) (*t*_5_ = −4.01, *p* = 0.003, Fig. 3A). Phenotypic changes were also more parallel (*θ* ± *SD*) on average (permutation test, *p <* 0.001) in hot lines (39.32 ± 19.16) than cold lines (67.42 ± 23.3), consistent with the hypothesis that high temperatures impose stronger selection (Fig 3B). Compared to Monte-Carlo simulations of random angles in 7-dimensional space, the mean pairwise angles for both cold and hot lines were more parallel than expected by chance (two-sided test, *p <* 0.05, 10^5^ iterations). Moreover, parallelism was greater within than between genetic backgrounds for both hot and cold lines (Table S1), suggesting a role of historical contingency in dictating the repeatability of phenotypic evolution.

**Figure 3:**
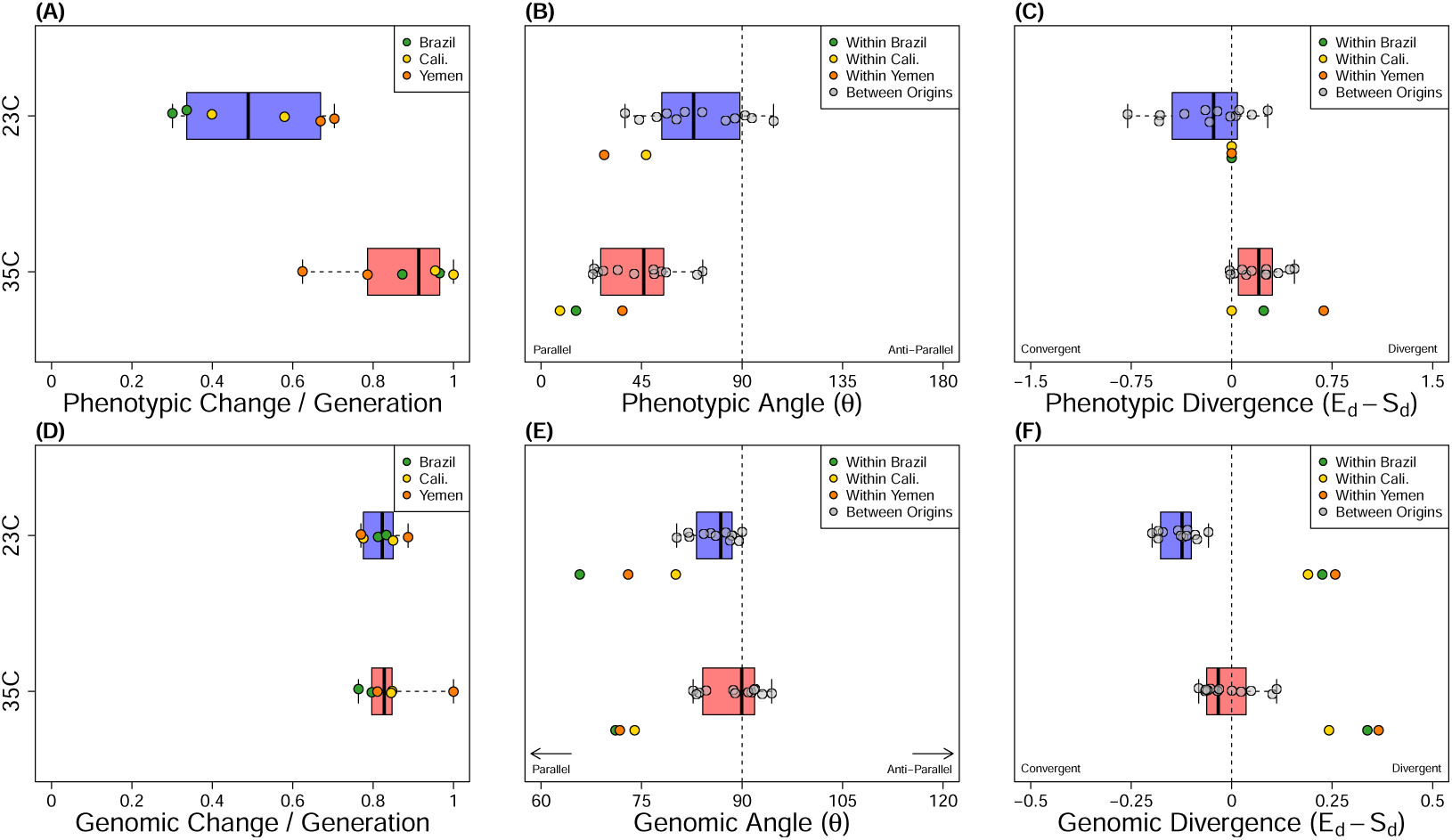
Distribution of all phenotypic and genomic evolutionary rates (A & D), pairwise angles (B & E) and measurements of convergent or divergent evolution (C & F) for cold (blue) and hot (red) lines . All genomic estimates are based on the set of putatively selected SNPs (n=475,194). Angles and divergences are given for pairwise comparisons between lines of different (grey points) and same (colored points) genetic background. Angles of 0° represent completely parallel responses, while angles of 90° represent uncorrelated responses. Divergence was calculated as the difference in Euclidean distance between population-pairs at the end (*E_d_*) and start (*S_d_*) of experimental evolution, and are divergent when the distance between end points is greater than the distance between start points (*E_d_* − *S_d_ >* 0), or convergent if the opposite is true (*E_d_ S_d_ <* 0). Evolutionary rates have been scaled by the maximum evolutionary rate in the dataset, while divergences have been scaled relative to the distance between the two most differentiated ancestors. Inestimable phenotypic angles due to sampling error are not shown (Within Brazil, 23°C).

Despite increased parallelism during adaptation to heat, cold lines generally exhibited less divergence (or greater convergence) relative to hot lines (Fig 3B) (*Div*_23_ = −0.14 ± 0.29; *Div*_35_ = 0.22 ± 0.20, permutation test, *p <* 0.001). While this result, to some extent, can be explained by the overall smaller magnitudes of per-generation phenotypic change in cold relative to hot lines, it also suggests that non-parallelism among cold lines in part resulted from convergent evolutionary trajectories of differentiated ancestral populations, making cold lineages become more similar to each other in some phenotypic traits during experimental evolution (Fig. 2A).

### The genetic basis of thermal adaptation is polygenic and governed by private alleles

We estimated allele frequency shifts and selection coefficients using wholegenome sequences (pool-seq) from all 12 evolved populations and their 6 ancestors (Fig. 1A). Consistent with stronger selection at warmer temperature, hot lines experienced greater rates of allele frequency change at candidate loci and were estimated to have lower *N_e_* than cold lines (Table S2). This pattern was similar for selection coefficients which take into account drift based on estimates of *N_e_*, with hot lines experiencing stronger average selection on all three backgrounds (Table S2).

Thermal adaptation was highly polygenic, involving several thousands of candidate SNPs (Table S3). Although it is unlikely that these SNPs are causal, several distinct peaks can be observed of SNPs deviating from drift expectations across almost all chromosomes in all populations (Fig. 2B, S2). We identified SNPs under putative selection on each genetic background separately and assigned these SNPs into four categories: Synergistically pleiotropic - SNPs selected in the same direction across both thermal regimes, antagonistically pleiotropic - SNPs selected in opposite directions across regimes, and private cold and private hot - SNPs selected in only the cold or hot regime (Fig. 4A). Contrary to typical theoretical assumptions, overall patterns indicate that thermal adaptation is not governed by antagonistic pleiotropy. Instead, thermal adaptation was primarily characterized by privately selected alleles. There were also more SNPs evolving in the same direction between temperature regimes than in opposite direction. These general patterns seem not affected by potential detection bias against pleiotropic, relative to private, SNPs (Fig. S3). Compared to forward in time simulations from ancestral populations (R package poolSeq v.0.3.5.9), there were more SNPs shared between the two line replicates within each background than expected under drift (all *p <* 0.001). However, most candidate SNPs were unique to a particular genetic background, with only private alleles selected at cold temperature showing modest repeatability across backgrounds (Fig S4).

**Figure 4:**
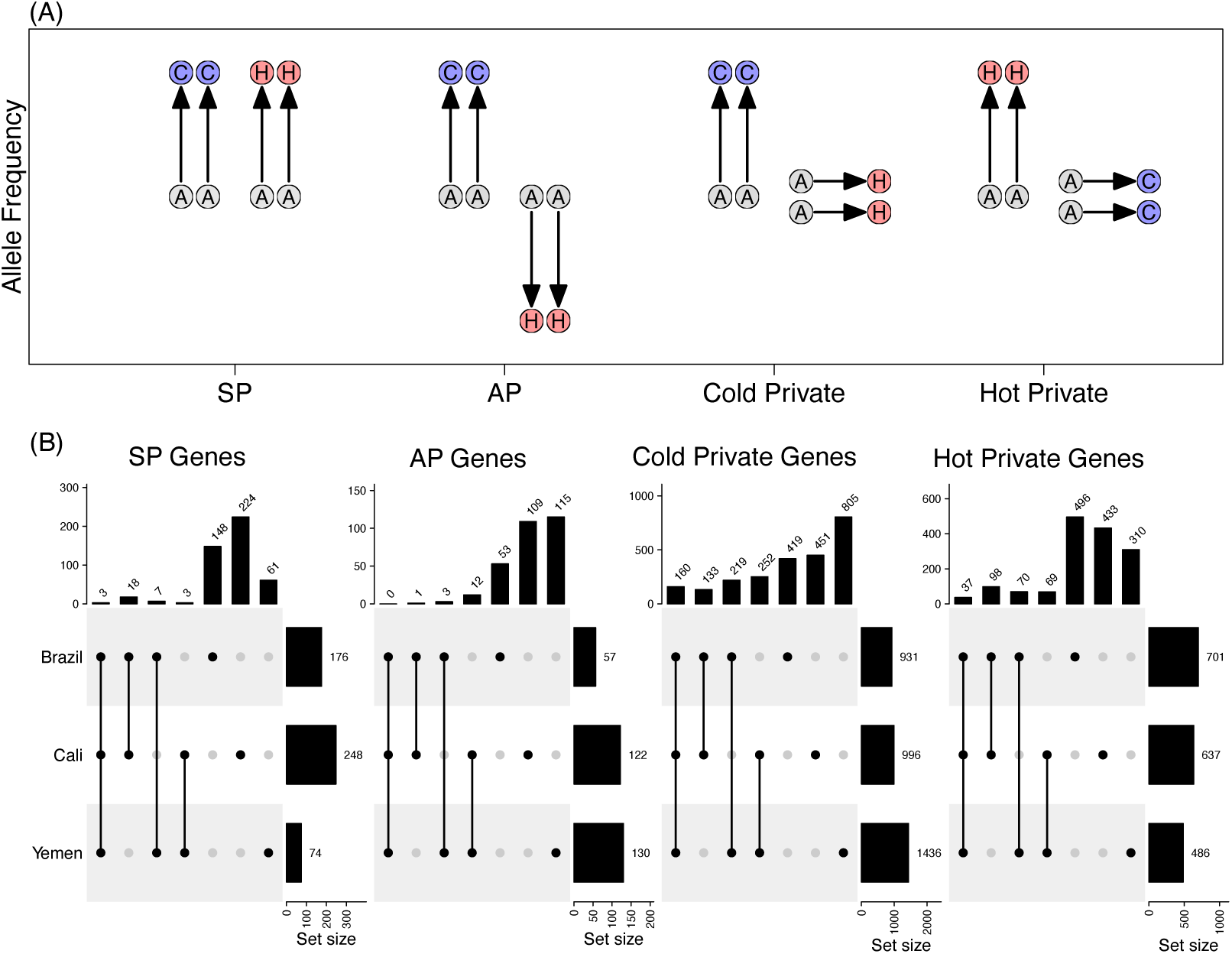
(A) Schematic showing how SNP and gene classes were assigned. For each class, we require that both replicates of a particular background behave similarly (i.e., more extreme than the 1st or 99th percentiles with the same direction of allele frequency change in both replicates). The four classes identified are: 1) Synergistically pleiotropic (SP) SNPs which are selected for in the same direction between regimes, 2) antagonistically pleiotropic (AP) SNPs which are selected in opposite directions between regimes, 3) private cold SNPs which are selected for only in the cold regime, and 4) private hot SNPs which are selected for only in the hot regime. (B) Upset plots of gene sets based on corresponding SNP set classes (see Fig. S4) located in protein-coding regions. There is significant excess of shared privately selection genes, and more gene sharing among genetic backgrounds adapting to cold temperature.

As expected, there was more repeatability at the level of genes (Fig 4B), with an excess of shared variants among all pairwise comparisons across origins (all *p <* 0.01). Across all six cold lines, there were 296 shared genes targeted by selection (i.e., containing a putatively selected SNP), which are many more than expected by chance (expected = 2.33, *p <* 0.001), while hot lines shared 51 genic targets (expected = 0.11, *p <* 0.001) (see Table S5 for associated GO terms). Repeatability was again more pronounced for privately selected genes and at cold temperature. Jaccard indices of the overlap of candidate genes indicated that cold lines show greater repeatability (0.32±0.06) compared to hot lines (0.20±0.06) (permutation test, *p <* 0.001) (Fig S5). We performed detailed scans across the genome for outlier windows showing recurrent allele frequency change among replicate lines using the R package AF-vapeR. This revealed that cold adaptation was characterized by regions which were selected upon across nearly all replicate lines (Fig. S6A), while heat adaptation was more often characterized by background-specific genomic responses (Fig. S6B). Hence, thermal adaptation seems best described by polygenic adaptation with effects private to each temperature, with modest repeatability across genetic backgrounds for heat-adaptation but higher repeatability when evolution proceeded at cold temperature (Fig 4B).

Although no antagonistically pleiotropic genes were shared between all backgrounds, such genes shared between two origins contained a cGMP-dependent protein kinase which is associated with thermotolerance in *D. melanogaster* larvae [40], and the gene *couch potato* (*cpo*) which is associated with diapause climate adaptation in both *D. melanogaster* [41] and *Culex pipiens* [42]. Gene set enrichment analysis (Fig S7) revealed that both synergistically and antagonistically pleiotropic genes were over-represented for processes such as oxidation-reduction processes, double-strand break repair, and DNA recombination, which may indicate general responses to suboptimal temperature [33]. The genetic backgrounds shared many more processes for privately selected gene sets. Notably, the GO term “Homophilic celladhesion via plasma membrane adhesion molecules (GO:0007156)” is found in almost all gene set categories, across all backgrounds. This GO term was also identified in several *Anolis* species in relation to adaptation of thermal niches [43] and cold tolerance in *D. melanogaster* [44]. As the mechanical and functional properties of the plasma membrane are greatly influenced by thermal fluctuations [45], it seems likely that this biological process is a strong candidate for shared targets of selection across thermal gradients.

### Genomic evolution is more predictable at cold temperature

We estimated the repeatability of genomic adaptations taking the same approach as for the phenotypic data and quantified angles between vectors of evolutionary change (here represented by shifts in allele frequencies) between lines deriving from the same or different genetic backgrounds. We aimed to ascribe observed temperature-specific patterns to three key factors expected to impact the predictability of evolution; i) environmental differences in the strength of selection (expecting more repeatability at hot temperature), ii) differences in available ancestral genetic polymorphisms at selected genes, and iii) genetic epistasis between selected genes and differentially fixed loci in the three ancestors. To achieve this, we first conducted analyses using all SNPs to detect candidate loci under selection on each of the three genetic backgrounds. We then repeated the analyses using only those SNPs that were shared among the backgrounds, expecting our estimates of repeatability to increase if differences in ancestral polymorphisms were key in driving the patterns.

Despite greater phenotypic parallelism in response to heat, hot lines did not exhibit greater parallelism at the genome level (*θ̄*_23_ = 83.42 ± 6.7, *θ̄*_35_ = 85.48 ± 7.8, permutation test, *p* = 0.15) (Fig 3C). Note that relatively orthogonal measures of parallelism compared to phenotypic estimates is ascribed to the difference in dimensionality of the two analyses. Indeed, the observed pair-wise angles among both cold- and heat-adapted lines were much smaller than expected by chance (Monte-Carlo simulations, *p <* 0.001, 10^5^ iterations; Fig. S8). Just as for the estimates of phenotypic repeatability, within-background comparisons were more parallel (*θ̄*_23_ = 72.96 ± 7.16; *θ̄*_35_ = 72.28 ± 1.5) than between-background comparisons, and especially so for heat-adaptation (*θ̄*_23_ = 86.03 ± 3.25; *θ̄*_35_ = 88.77 ± 4.19), suggesting an important role of historical contingency in dictating genomic repeatability (Table S4). We note that these qualitative patterns hold true even when estimating parallelism from temperature-specific sets of SNPs with differing dimensionality and are unlikely to be due to differences in the number of selected sites between regimes (Fig. S11C-D, S8A-B).

Cold lines tended to diverge less (or converge more) in genomic space than hot lines (permutation test, *p* = 0.011). This pattern is true both within (*Div*_23_ = 0.22 ± 0.03; *Div*_35_ = 0.32 ± 0.06) and between backgrounds (*Div*_23_ = −0.13 ± 0.04; *Div*_35_ = −0.01 ± 0.06) (Fig 3F). Unlike phenotypic rates, genomic rates of change at selected loci were not different between hot and cold lines (*t*_5_ = −0.59, *p* = 0.57 (Fig. 3D).

That we see greater phenotypic repeatability for hot lines, but higher genomic repeatability across cold lines for all genomic metrics is counterintuitive and opposite of our expectations. We propose three non-mutually exclusive mechanisms that may contribute to such patterns. First, due to stronger selection at high temperature, hot lines also experience more genetic drift, as evidenced by lower *Ne*. This could contribute to lower predictability of genomic changes across lines. However, *Ne* was not strongly correlated with the repeatability of observed evolutionary trajectories (Fig. S9A–B) nor with divergence at candidate loci within regimes (Fig. S9C–D).

Second, differences in ancestral polymorphism at selected loci may contribute to historical contingencies and low repeatability across geographically separated populations [23]. To test this hypothesis, we re-estimated all measures of genomic repeatability based on only those putatively selected SNPs that segregated in all ancestral populations (n=119,630, or ∼25% of the original set), expecting to see significantly higher repeatability, especially for heat-adapted lines, if differences in ancestral polymorphism have contributed to the observed temperature-specific patterns. However, although genomic repeatability was slightly increased overall for shared polymorphic sites, the greater effect of genetic background on parallelism and divergence observed for heat- relative to cold-adaptation remained (Fig. S10–S11; Table S6). Additionally, when comparing the overlap between backgrounds of candidate genes in the privately selected gene sets (which contain the majority of candidate SNP), we found no difference with respect to using all SNPs or only those shared among all backgrounds (Fig S12, all Chi-square test *p >* 0.05). Hence, differences in shared ancestral polymorphisms seem to have played a minor role in shaping the observed genomic repeatability between temperatures.

Third, it is well recognized that gene interactions (epistasis) can contribute to unpredictable evolutionary outcomes [46], and that such epistasis might depend on the environment [27, 47]. Indeed, epistasis is predicted to be a natural property of metabolic networks, and as all metabolic processes are strongly temperature-dependent, it is possible that epistasis between differentially fixed variants in the three genetic backgrounds and selected variants may have been stronger at hot temperature. In Fig 5 and Supplementary (Fig. S18), we illustrate that our temperature-specific estimates of genomic adaptation (i.e., excess of privately selected SNPs) and repeatability (lower at hot temperature) are consistent with predictions from a simple model of thermal adaptation assuming exponential increases in both enzyme reaction rates (Fig. 5A) and molecular failure rates (e.g. loss of enzyme stability; Fig. 5B) with increasing temperature [37, 48]. This heuristic model has been used previously to argue that hot temperature results in increased epistasis by causing proteins to evolve marginal thermostability, such that further mutational or thermal perturbations can be either inconsequential or highly deleterious depending on the wildtype protein [37, 38, 49–51]. Nevertheless, the mechanistic basis underlying the observed temperature-specificity of evolutionary repeatability needs to be investigated further.

**Figure 5:**
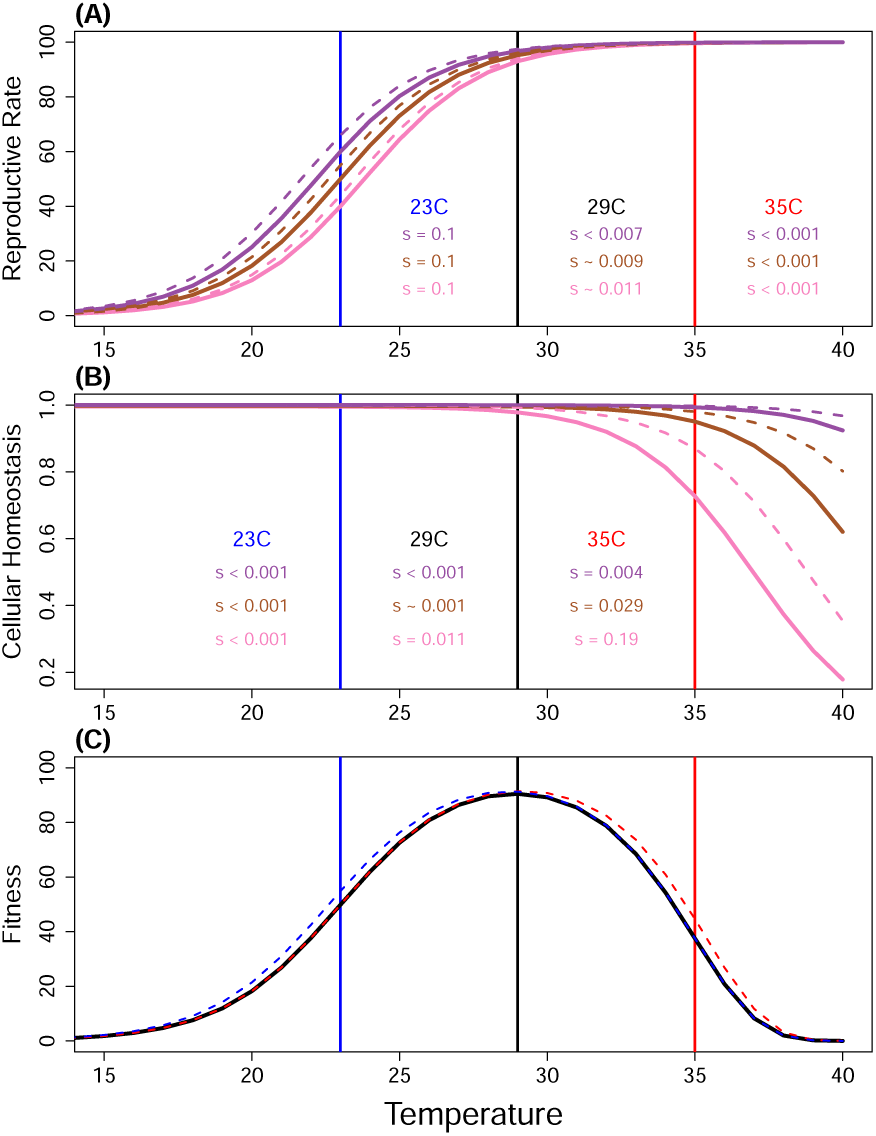
A prediction of temperature-dependent epistasis for fitness based on thermodynamics of cellular processes. For each plot, colored vertical lines denote the temperature of our thermal regimes (23°C and 35°C). **A)** Reproductive rate increases exponentially with temperature at the colder range due to relaxation of thermodynamics constraints on enzymatic reactions, but starts to follow a pattern of diminishing returns at warmer temperatures due to ecological constraints placing limits on reproductive output. Illustrated for three hypothetical genetic backgrounds (purple, brown and pink) with wildtypes (solid lines) and a mutant (broken lines) with a 10% increase in enzymatic reaction rate at 23°C. This results in strong selection on the mutation at cold temperature for all backgrounds as these are far from their optimal reproductive rate, but weak selection at hot temperature at which all backgrounds are close to the maximal achievable reproductive rate determined by ecological factors. Epistasis is weak and selection is similar across backgrounds at all temperatures. **B)** Warm temperatures increase a range of molecular failure rates, which results in less enzyme ready to catalyze reactions, and more misfolded proteins within cells, leading to protein toxicity and depressed fitness. Illustrated for three genetic backgrounds with stable (green), intermediate (orange) and unstable (yellow) wildtype protein. A mutation increasing stability (broken lines) was introduced on each background. Selection on the mutation is weak at cold temperature for all backgrounds but can become very strong at hot temperatures depending on the wildtype protein stability, and strong epistasis for fitness results. **C)** Fitness, as the product of reproductive rate and molecular failure rate, is depicted for the wildtype (black), and the mutants with increased enzymatic reaction rate (blue) and increased protein stability (red) for the pink background.

### Does Genomic Evolution Predict Phenotypic (mal)adaptation?

To assess the premise of predicting phenotypic adaptation from genomic data alone, we calculated “offsets” for both types of data, describing the level of genetic and phenotypic maladaptation relative to the most well-adapted line which we used as the reference [52, 53]. We performed this calculation for each of the three backgrounds and for hot and cold assay temperature separately. We here focus on offsets at hot assay temperature as we see more clear signs of local adaptation to heat than cold, and because phenotypic estimates of cold-adaptation were smaller and many times not statistically significant (Fig. S1). For completeness, we present the results for cold assay temperature in (Fig. S13 and S14).

Genomic offsets were strongly predictive of phenotypic distance and fitness offsets among populations reared at hot temperature for lines of the same geographic origin as the reference (*r̄_P heno_* = 0.88, ANCOVA *F*_1,9_ = 13.6, *p* = 0.005; *r̄_F it_* = −0.74, ANCOVA *F*_1,9_ = 7.23, *p* = 0.025). However, when applied to lines of different geographic origin, the predictive power of genomic data was severely reduced (*r̄_P heno_* = 0.22, ANCOVA *F*_1,30_ = 0.86, *p* = 0.36; *r̄_F it_*= −0.29, ANCOVA *F*_1,30_ = 1.24, *p* = 0.27) (Fig. 6). These results remained qualitatively unchanged when considering only SNPs which were polymorphic in all ancestors (S15), suggesting that differences in ancestral polymorphisms were not *per se* causing the poor predictability across genetic backgrounds. . Hence, selected sites identified in a particular origin do not contribute greatly to phenotypic or fitness variation in other origins, which is congruent with estimates of both phenotypic and genomic repeatability being greater within than between origins (Fig. 3).

**Figure 6:**
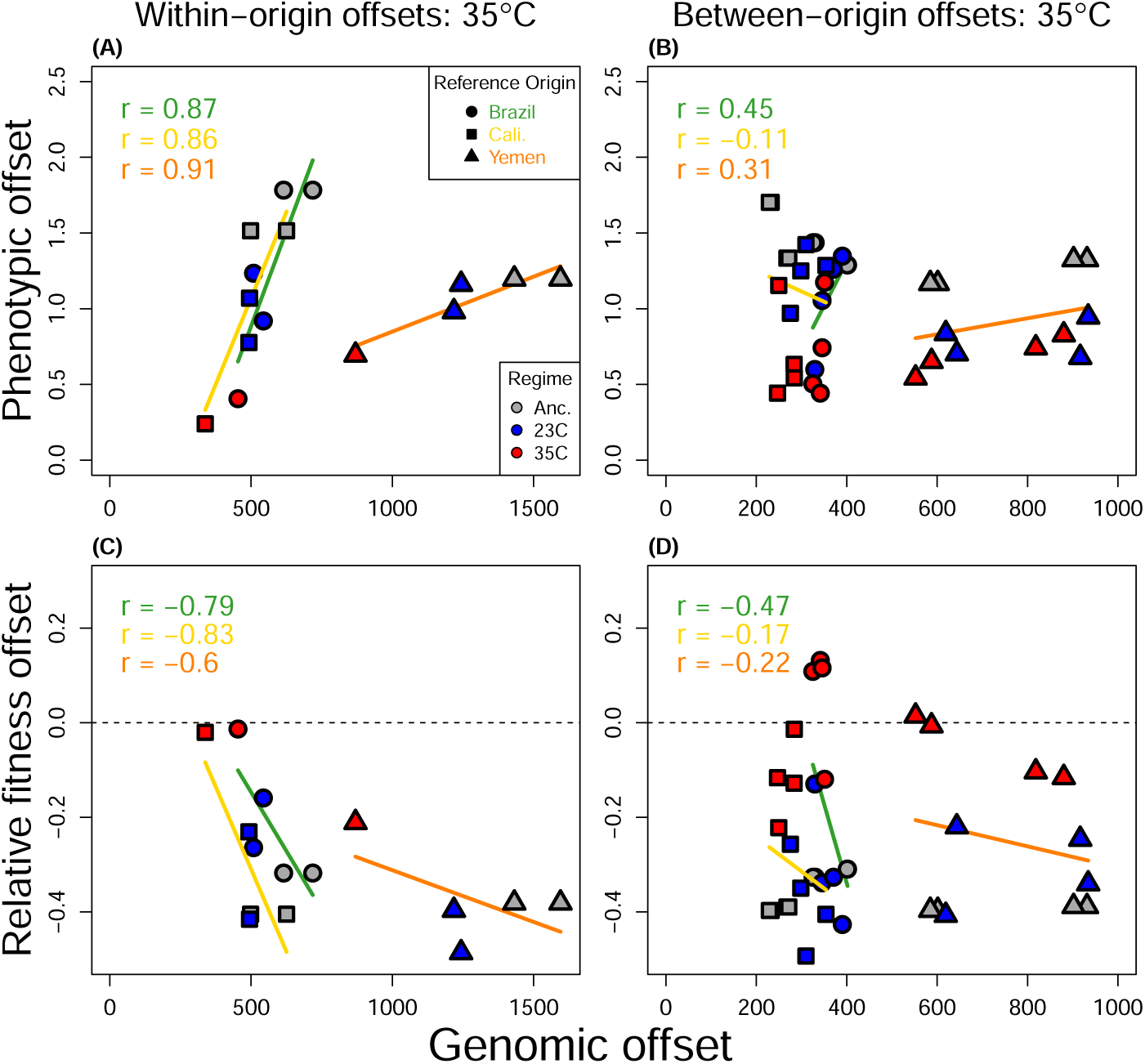
Genomic predictors of phenotypic divergence and laboratory fitness at hot temperature (35°C). Genomic offsets were calculated per origin based on SNPs whose p-values for allele frequency change fell within the top 0.001th quantile (n=10546–10649). Phenotypic offsets were also calculated per origin. Laboratory fitness offsets were calculated as a line’s lifetime adult offspring production/generation time, relative to that of the line with the highest fitness, which was designated as the reference population. Similarly, phenotypic divergence offsets were calculated as the Euclidean distance in scaled trait-space to this reference population. Offsets are organized by predictions within genetic backgrounds (A,C) and between genetic backgrounds (B,D). Correlations and regression lines are colored by the origin of the reference population, while symbol colors refer to ancestral (grey), cold (blue) and hot (red) lines. Genomic offsets predict maladaptation within (A, C), but not between (B, D), geographic origins.

Interestingly, the genomic offsets were typically larger between lines of the same genetic background compared to offsets between backgrounds. This suggests that, while selected sites identified in a given reference line contribute to fitness/phenotypic variation among lines of the same geographic origin, these sites may be mostly fixed (in the direction of the reference line) in lines of the other origins. It could be, then, that by focusing on SNPs which show signs of selection in a limited set of study populations, the importance of already-fixed sites that cannot contribute towards genomic offsets may be greatly underestimated in genomic prediction of geographically distant populations. To explore if this possibility contributed to the poor genomic predictions of phenotypic adaptation applied across origins, we estimated the correlation between fitness-offsets and genomic offsets basing the latter on all candidate SNPs per thermal regime (i.e. identified in any of the six line replicates), with the reasoning that this should better capture and include differentially fixed sites. However, even when all putatively selected sites were considered, predictive power remained high in the case of within-origin estimates (*r* = −0.87), and low for between-origin estimates (*r* = 0.32) (Fig. S16B). Compared to offsets utilizing selected SNPs, sets of randomly chosen SNPs performed worse for within-origin estimates, but on-par with across origin predictions (Fig. S17).

## Discussion

Climate change is disrupting ecological niches and changing species-distribution patterns across the globe [54, 55]. For many taxa, the potential to genetically adapt to these changes is necessary to avoid extinction. Approaches harnessing genomic data to predict adaptive potential in key ecological traits of threatened populations may provide means to focus conservation efforts and are becoming more common, but their limitations are debated [15, 17, 18, 56, 57]. Here, we adapted seed beetles from three different geographic origins to cold and hot temperature and assessed both life-history and genomic adaptation to 1) determine the genetic basis of thermal adaptation, 2) quantify the repeatability of evolution across biological levels of organization, and 3) evaluate the prospects of genomic prediction of adaptive potential across geographically isolated populations. We show that adaptation to both cold and hot temperature is highly polygenic. Evolutionary responses were more repeatable within, than between, genetic backgrounds, demonstrating an important role of historical contingency in dictating evolutionary predictability. Heat adaptation resulted in greater phenotypic rates of change, indicative of high evolutionary potential under future climate warming. However, despite a high repeatability of phenotypic evolution at hot temperature, adaptation at the genomic level was more repeatable at cold temperature, with the effect of genetic background playing a larger role for heat-adaptation. Indeed, while genomic offsets for heat-adaptation predicted maladaptation in lines from the same geographic origin reasonably well, they failed to predict maladaptation in lines from different origins. This suggests that the prospects of using data on genomic-phenotype correlations to predict more broad-scale geographic patterns of (mal)adaptation and evolutionary potential under future climate warming may be limited. Complementary hypotheses exist to explain the observed patterns, which we discuss below.

### Genetic basis of thermal adaptation

Thermal adaptation was found to be highly polygenic across all lines and regimes, with several thousand polymorphisms under putative selection, which is predicted to result in high genetic redundancy and low repeatability of adaptation at the genomic level [13, 58]. Our results indicate that thermal adaptation is largely characterized by alleles with effects limited to a specific thermal range, suggesting that evolutionary potential under climate change may be fueled by alleles with conditional fitness effects that are relatively abundant at ancestral and benign thermal conditions but become exposed to selection at thermal extremes. This notion also corresponds well with predictions of temperature-dependent selection on enzymes based on the thermodynamic basis of protein function [37, 59, 60] (and Fig 5). Our results also mirror experimental evolution studies in both *Drosophila melanogaster* [61] and *D. simulans* [25, 62] in which adaptation was polygenic and few targets of selection overlapped between opposing thermal selection regimes.

A possible exception to this main pattern is the observed divergence along one genomic dimension, explaining roughly 6% of all the variation at selected sites, that occurred in opposing directions for hot and cold lines (Fig. 1E, genomic PC3). This is in contradiction to our analyses of single SNPs and genes, where very few sites showed signs of antagonistic pleiotropy. It is well recognized that polygenic adaptation can result in genetic redundancy with stochastic and modest allele frequency changes at single sites, where allelic effects often show epistasis for fitness during the process of adaptation, causing a highly unpredictable genomic signal at the level of single SNPs [13, 63]. While this points to the general problem of detecting polygenic selection at weakly selected loci, our results also highlight that overall genomic signals, such as that depicted along PC3, can still be picked up and be informative of the basis of adaptation, given sufficient experimental replication.

### (Non)parallel adaptation to thermal stress

Evolutionary trajectories were generally more parallel for comparisons within, than between, backgrounds for both phenotypic and genomic adaptation, demonstrating an important role of historical contingencies in shaping evolutionary outcomes. Similar results were found in studies of temperature adaptation in different founders of *Drosophila simulans* [25, 62]. Likewise, studies of wild populations have found significant, but modest overlap between climate adaptation loci among independently evolving lineages [64–67].

Biophysical models of protein stability predict that higher temperatures should generate stronger purifying selection [37, 38, 60] and increased positive selection on alleles that buffer thermal stress [33, 68]. We therefore expected both faster and more repeatable adaptive evolution at hot temperature. We found evidence at the genomic level that warm temperature indeed imposed stronger directional selection based on estimated selection coefficients (Table S2). However, contrary to our expectations, we observed that heat-adaptation was only more parallel for phenotypes. Our follow-up analyses did not provide evidence for genetic drift being a causal explanation for the lack of genomic repeatability. Likewise, our estimates of repeatability based only on those SNPs that were shared between all ancestral populations suggest that the observed patterns are unlikely to be solely explained by differences in the access to (adaptive) standing genetic variation [23]. As a final explanation, epistasis between selected SNPs and differentially fixed variants in the three ancestors may have been magnified at hot temperature. Epistatic interactions between different proteins or between different SNPs within the same protein coding sequence are ubiquitous [27, 69, 70], and temperature-dependent increases in such interactions are emergent properties of the assumptions in simple biophysical models of protein thermodynamics and function [37, 51, 60] (Fig. 5) and have been observed in empirical studies [71]. Although we cannot explicitly test this hypothesis with the current data, the low explanatory power of both genetic drift and differences in ancestral polymorphism, together with the larger difference in repeatability between the hot and cold regime observed across (compared to within) genetic backgrounds, suggest that temperature-dependent epistasis may be causally linked to the lower genomic repeatability observed at hot temperature. This suggests, somewhat paradoxically, that the same thermodynamic mechanism that causes hot temperatures to impose strong selection, resulting in repeatability of adaptive outcomes at the phenotype level, may at the same time make genomic evolution more unpredictable by increasing epistatic interactions.

### Predicting (mal)adaptation from genomic data

Leveraging genomic datasets to predict the fates of threatened populations is an important aim in ecological and conservation genetics. However, both genetic redundancy [25] and epistasis [72] obfuscate the genotype-to-phenotype map, potentially reducing the power of using genomic data to make inferences about adaptive potential. In accordance with this notion, our genomic offsets were predictive of phenotypic adaptation to heat when applied within a single genetic background, but poor predictors when applied across genetic backgrounds. This suggests that predictions of adaptive potential under future climate change may be problematic when applied on large geographic scales, maybe to the extent that standard estimates of (non-specific) genomewide genetic diversity may offer similar accuracy in predictions. Indeed, both this study (Fig. S17) and a comparison of several methods for conducting genomic offsets found that sets of randomly selected SNPs can sometimes better predict (mal)adaptation than identified candidate SNPs when applied across strongly differentiated populations [73].

A possible explanation for low predictability between genomic and phenotypic data is that many changes in allele frequency may not be causal for changes in phenotype. Our seed beetle populations experienced low effective population sizes (*N̄_e_* = 215.4) and strong selection, which likely resulted in relatively large allele frequency changes due to genetic drift and draft, which may have resulted in skewed associations between genomic and phenotypic change. Moreover, the relatively low census population sizes (300- 400 individuals) during laboratory maintenance will have contributed to further fixation of alternative alleles in the three ancestral populations, amplifying differences between the ancestors. Nevertheless, the population sizes studied here are probably representative of many threatened species, which are the prime targets for genomic prediction. Moreover, our conclusions on temperature-specific repeatability of genomic changes within and between genetic background are unlikely to be affected by low *Ne*, as our analyses focused on alleles under strong selection (s¿0.01), and remained qualitatively unchanged when only considering genetic variants shared among all three backgrounds.

The particular metrics used for phenotypic and performance measures should also be given consideration [52, 53]. While whole-genome analyses of predictability can typically incorporate all individual ‘units’ which compose the response to selection (i.e., changes in allele frequency across the genome), phenotypic or fitness analyses can be sensitive to the set of traits measured [74]. Predictability should be highest at the level of fitness, relative to underlying traits [75]. The suite of traits we measured is expected to by highly associated with fitness during temperature adaptation, and should thus exhibit high predictability in evolutionary changes. Some traits chosen here have also shown strong parallel responses in *Drosophila* [25]. Nevertheless, we note that our assays did not include juvenile competition traits, that may have been central in driving cold-adaptation [76], potentially explaining the lack of identified local adaptation and poor correlations between genomic and fitness offsets at 23°C.

## Conclusions

Our work combines evolutionary change at the phenotypic and genomic level, something so far rarely done in thermal adaptation studies [31, 32, 62, 77]. While phenotypic evolution was more rapid and repeatable at warm temperature, the opposite was true for genomic change, with historical contingencies obfuscating the relationship between genomic and phenotypic adaptation to heat. Multiple non-mutually exclusive mechanisms may have contributed to this pattern. Importantly, theory predicts that many of the same mechanisms that should increase predictability at high temperatures (e.g., strong selection on cellular homeostasis) may contrarily result in reduced genomic predictability due to the biophysical properties of cells, resulting in epistasis, and the demographic properties of populations, resulting in genetic drift/draft. Our findings suggest that predicting adaptation to future warming climates using genomic data may become increasingly difficult, especially when extrapolating across geographically isolated populations, thus placing emphasis on sampling many populations to more confidently describe hypotheses regarding broad patterns of evolutionary repeatability [78]. For genomic predictions, there is a scarcity of experimental evolution studies to validate and refine predictive methods, especially with regards to the role of epistasis in dictating the relevance of genomic predictions across larger geographic scales. Such studies will be invaluable for advancing predictive accuracy of forecasts of threatened species.

## Materials and Methods

### Study Species

The seed beetle, *Callosobruchus maculatus*, infests human stores of grain legumes. Females attach eggs to the surface of seeds and larvae burrow into and develop within a single seed. Adults then emerge and reproduce, typically within 3-4 weeks, without the need for additional nutritional resources and a full life cycle can be completed within less than a month at benign temperatures; egg-to-adult development time is thus a good approximation of generation time in laboratory conditions. Because *C. maculatus* has been associated with stored legumes for thousands of years, laboratory conditions are a good approximation of its “natural” environment [79]. The experimental lines used for this study are described in detail in [80] and [39]. Briefly, outbred stocks were initially collected from three geographic populations (Brazil, California, and Yemen) and kept in laboratory conditions for *>*60 generations. The mitochondrial haplotypes of each population were introgressed in an orthogonal fashion into each nuclear background through repeated backcrossings over 16 generations, such that each mitochondrial haplotype existed in equal frequency on each nuclear background [81, 82]. The introgressed lines were kept as separate populations for ca. 100 generations, at which point females from each line were backcrossed to males from the source population with the same nuclear background to refresh nuclear genetic diversity. These newly backcrossed lines were used as the three base populations (Brazil, California, and Yemen) for our experimental evolution. At the time of study, these stocks had been reared on black-eyed beans (*Vigna unguiculata*) under standard laboratory rearing conditions of 29°C in 12:12 light/dark cycles and 50% relative humidity, and *N* = 300 − 400, for *>*200 generations.

### Ancestral Thermal Performance

To assess the temperature tolerance and preference of the three ancestors, we measured how laboratory fitness changed with rearing temperature. Laboratory fitness was defined as the number of adult offspring produced per mating couple, divided by the egg-to-adult development time of those offspring as an estimate of generation time; this measure should thus closely corresponds to each population’s predicted intrinsic population growth rate. Each population was allowed to lay F1 eggs on the standard host, black eyed beans (*Vigna unguiculata*) at 29°C. The beans were then split among 23, 29 and 35°C. Once F1 adults emerged, 20-30 mating couples per temperature were isolated and allowed to mate and lay eggs for their entire lifespan. Once emerged, F2 adult offspring were recorded for their development time, frozen, and later counted. To also get estimates of laboratory fitness at more extreme temperatures, 10 F1 mating couples emerging from 35 and 23°C were moved to 37 and 17°C, respectively, and assayed. We note here that, because these F1 parents were not developing at the extreme temperatures, effects on their fertility from developing at the temperature extremes are not accounted for. This may have underestimated the stress imposed by 37°C presented in Figure 1. Thermal performance curves presented in Figure 1 were fit using a third-degree polynomial using the “spline” function in R. When doing so, we fixed laboratory fitness at 38°C to zero, as we have not been able to propagate any populations of *C. maculatus* at this temperature [38, 83].

### Experimental Evolution

Each replicate line was maintained on 250 ml black eyed beans in a 1 liter glass jar kept in a climate cabinet set at 50% relative humidity at the given evolution regime temperature (23°C or 35°C) (Fig. 1A). Once adults emerged from beans, 600 beetles were transferred to a new jar with fresh beans and allowed to mate and then lay eggs until their death. Beetles were transferred at the peak of hatching, so as not to apply strong direct selection for faster development time. See [80] and [39] for further details.

### Phenotypic data and analysis

We re-analyzed previously collected data on thermal adaptation in seven lifehistory traits of females [39], including three core traits: lifetime offspring production (LRS), adult weight (AW) and juvenile development time (DT) (Fig. 2A), as well as four rate-dependent traits measured over the first 24 hours of female reproductive lifespan: weight loss (WL), water loss (WVP), early fecundity (eggs laid over the first day of adult female reproduction) and mass-specific metabolic rate (ml CO_2_/min/mg). As reported by [39], Hot-adapted lines have evolved greater body mass, weight loss, and reproductive output than cold lines, whereas rate-dependent traits show signs of local adaptation to temperature signified by statistically significant regime-by-assay temperature interactions.

These data were collected at generation 45 for cold-adapted lines and 60 for hot-adapted lines, but at the same time in a large common garden experiment removing (non-genetic) parental effects and including the two assay temperatures corresponding to the experimental evolution regimes (23°C and 35°C). Phenotypic data from the ancestral populations were not collected at the time of line establishment, but were scored in a separate experiment. Because each line had been lab-adapted for *>*200 generations, we assume that minimal lab adaptation had occurred after thermal regime establishments, and that these phenotypic data approximate the phenotypes of the founding populations. To estimate any block effects due to differing sampling times, a reference line was used in both the experiment on ancestors and E&R lines, and the relative difference in trait means of this reference line in the two experiments was used to standardize the ancestral data to minimize the possibility that block effects were wrongfully assigned as evolved differences between ancestors and evolved lines. We note that the standardization was made for each trait by averaging over temperatures, and thus, does not contribute to the differences in the repeatability and magnitude of phenotypic adaptation between the temperatures and genetic backgrounds that we study here. Data on LRS was then further gathered in a large second common garden including both ancestors and evolved lines, after 80 (cold), 120 (hot) and 125 (ancestors) generations of evolution. Methods for data collection are thoroughly described in [39].

By including ancestral data, we were able to assess the direction and magnitude of phenotypic evolution. Following [84], we define parallel evolution here in a geometric sense by the angles of phenotypic change vectors between evolving populations. Although it should be emphasized that (non)parallelism is a continuous measure, three general categories can be made of the degree of (non)parallelism. In multi-dimensional trait space, evolution is parallel when the vectors of phenotypic change between evolving populations result in small angles (*θ <* 90°), and anti-parallel when angles are large (*θ >* 90°). Evolution can also result in orthogonal (i.e. uncorrelated) changes when *θ* ≈ 90°. Following [74], we calculated the 6 × 6 inter-population correlation matrix of evolutionary change vectors for each regime, and subsequently arccosine transformed correlations to angles. Because change vectors for each of the two replicate lines of a given origin were calculated using the phenotypic scores of the same ancestral population, shared measurement error in the ancestor will have contributed to more similarity between line replicates of the same origin, relative to similarity in phenotypic change vectors of lines of different origin (with separate ancestors used in calculations of change vectors). The angle between two change vectors can be decomposed into its components; 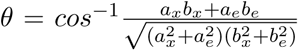 where *a_x_* and *b_x_* represent the vectors of population mean trait change for evolved lines *a* and *b*, and *a_e_* and *b_e_* represent the vectors of standard errors for each trait in the ancestors. This latter error component of the dot product (numerator) sums to zero when measurement errors are uncorrelated, such as the case when using different ancestors in calculations, but equals the squared standard error when using the same ancestor. To produce unbiased comparisons within and between backgrounds we therefore corrected for this contribution of shared measurement error by subtracting this error component from the within estimates of change vectors. For the two cold-adapted Brazil populations, this error component was larger than the observed mean trait change. The angle of phenotypic change between these populations was therefore not estimated due to the statistical uncertainty.

By comparing the phenotypic distances between populations at the start (*S_d_*) of experimental evolution (i.e. between ancestors) to the distance at the end (*E_d_*) (i.e. between the evolved lines), we estimated whether populations were diverging or converging phenotypically over the course of the experiment. These pairwise distances were calculated based on all 7 phenotypic traits. When evolved lines are more distant than the ancestors in trait-space (*E_d_* − *S_d_ >* 0), populations are diverging, while the opposite (*E_d_* − *S_d_ <* 0) is indicative of convergence. To estimate evolutionary responses to heat, we compared the Hot lines to their ancestors when reared at 35°C. Similarly, cold-adaptation was estimated by comparing Cold lines to their ancestors when reared at 23°C. To perform the evolutionary change vector analysis, we first mean-scaled all traits into unitless measurements for comparison with each other by dividing ancestral and evolved estimates by the grand-mean across all ancestors measured at the same assay temperature. Hence, rates of evolution are reported as proportional changes relative to ancestral trait means (see [85, 86]). We then calculated the vector of phenotypic change from the ancestral to evolved samples in the 7-dimensional trait-space. Similarly to the error correction conducted for angles, divergence within backgrounds will be overestimated relative to that between backgrounds due to the use of a single shared ancestral population in the former. We therefore corrected within-background distances by subtracting the squared standard errors of distance estimates in the two evolved replicates as 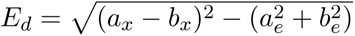, where *a_x_* and *b_x_* represent vectors of trait means, and *a_e_* and *b_e_* represent the vectors of standard errors, in evolved lines. For instances in which the error component was larger than the observed mean trait change component, the distances were set to zero (in all within-origin comparisons at 23°C and between the two replicate California lines at 35°C).

### Genomic data and analysis

For evolved populations, we isolated genomic DNA from pools of 60 male adults between generations 57-59 for cold lines and generations 67-69 for hot lines. Within each thermal regime, lines were sampled at the same time but varied in the number of generations of adaptation due to differences in development rate (Fig. 1A). Due to a paucity of beetles remaining after line-establishment, we were not able to sample the direct establishing ancestor. Instead, the third generation of heat-adapting lines was sampled and sequenced as a proxy of the ancestors (here simply referred to as ‘ancestors’ when considering genomic data); hence comparisons between evolved and ancestral lines represent 60-62 and 64-66 generations of divergence for the cold and hot regime, respectively. Adult beetles were collected and stored at -80°C. We purified DNA from the beetles using the King Fisher Cell and Tissue DNA Kit (Thermo Fisher Scientific). DNA fragment libraries for whole-genome sequencing (WGS) were generated at the Science for Life Laboratory (Stockholm, Sweden) from each pool by following the standard Illumina TrueSeq PCR-free library prep protocol in which 350bp fragments were retained. Purified genomic libraries were then sequenced on the Illumina NovaSeq 6000 platform. In total, two lanes of 150bp paired-end reads were generated.

We used BWA mem v.0.7.9 [87] to align the ∼5.8 billion ∼150-bp DNA sequences to a genome assembly for *C. maculatus* (accession: PRJEB60338). After sorting, merging, and de-duplicating (Picard Tools v.2.23.4, Broad Institute) our alignments, we then identified SNPs using the GATK best practices recommendations [88]. Variants were initially filtered based on GATK hard-filtering recommendations after manual and visual inspection of the data with the following parameters: *QUAL <* 20, *QD <* 2, *FS >* 20, *SOR >* 5, *MQ <* 40, −5 *> MQRankSum >* 5, −5 *> ReadPosRankSum >* 5. A second round of filtering was performed on regions of the genome that do not uniquely map to itself using GenMap v.2.4.1 (with parameters K31,E2) [89], sites with missing data across any sample, or *DP <* 20 across any sample. We retained 10,611,665 variants after filtering (∼10 variants per kb).

To measure the strength of directional selection, we first identified candidate loci consisting of putatively selected SNPs. To determine whether a SNP is likely under selection, we incorporated temporal information of allele frequency change from the ancestor to evolved hot and cold lines. We estimated the variance effective population size (*N_e_*) during the experiment for every replicate from patterns of allele frequency change from the total set of ∼10 million SNPs using the R package poolSeq [90]. Given the estimated values of *N_e_*, we then parameterized and tested a null model of evolution by genetic drift to determine the probability of observing the magnitude of allele frequency change for each SNP in each population. We calculated the probability of the observed allele frequency change from the ancestral to evolved populations based on a beta approximation to the basic Wright–Fisher model [91]. Following [92], we assumed *p_t_*|*p*_0_ ∼ beta(*α* + 0.001*, β* + 0.001), where 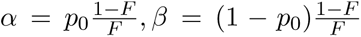, *p*_0_ and *p_t_* are the allele frequencies at the beginning and end of the interval, 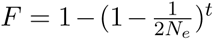, *t* is the number of generations between samples. We retained SNPs with allele frequency changes more extreme than the 0.01th or 99.99th percentiles of the null distribution per SNP in any population (Fig. 2B), resulting in a total set of 475,194 putatively selected sites among all replicates. Estimation of selection coefficients was performed with CLEAR [93]. We generated the input for CLEAR from the set of putative SNPs with scripts from the PoPoolation2 pipeline [94]. Estimations of *N_e_*, null model drift estimates, and selection coefficients all require information on the number of generations elapsed between timepoints; for each replicate, we therefore added or subtracted 3 generations for cold- and hot-adapted lines, respectively, to the generation parameter to account for drift that occurred between ancestral lines and our proxies of the ancestors.

Repeatability at the SNP level was measured, similarly to the phenotypic data, by geometric estimates of *θ* and divergence (*E_d_* − *S_d_*) of allele frequency change vectors between populations of the set of putatively selected sites. Note that because two replicate ancestral samples were available per background, error corrections were not implemented. Because we are interested in parallelism due to selection, we focused on the repeatability of evolutionary change occurring in the set of putatively selected SNPs.

We additionally scanned for genomic regions which may exhibit parallel or anti-parallel responses between replicate lines per regime with the software AF-vapeR (v.0.2.1) [95] implemented in R. We followed standard methods, calculating allele frequency vectors in windows of 200 SNPs each. Null expectations of allele frequency change were conducted by permuting allele frequency changes among populations. A total of 10,000 permutations were conducted across all chromosomes, with the number of permutations per chromosome proportional to the size of the chromosome.

### Repeatability and historical contingency for different classes of alleles

To explore if certain types of variants are more likely to contribute to repeated instances of thermal adaptation, we first identified SNPs which were assigned as being under selection in both replicate lines deriving from a particular background and regime. Because of this stricter criterion, we relaxed our null drift expectations to include SNPs with allele frequency changes more extreme than the 1st or 99th percentiles of the null distribution, resulting in 271,515–617,395 candidate SNPs being selected. For each genetic background and evolution regime separately, SNPs were then assigned into the four categories based on the patterns of evolutionary change observed between regimes (synergistically pleiotropic, antagonistically pleiotropic, private cold, and private hot) (Fig. 4A). We then compared the relative abundance of SNPs that showed overlap between the three genetic backgrounds for each category. The same analysis was carried out at the gene-level based on if the SNP fell within the protein coding region of a gene. The biological processes for each gene set were analyzed for enrichment using the R package topGO (v.2.38.1) and weight01 algorithm.

For each SNP category, we tested whether there was more overlap between backgrounds than expected by chance by performing forward in time drift simulations from all ancestral populations (R package poolSeq v.0.3.5.9). During each simulation iteration (n=1,000), 10,000 random SNPs were sampled to parameterize all lines. Each line was parameterized with the ancestral allele frequencies, the number of generations evolved, and the effective population sizes of respective line.

### Genomic and phenotypic offsets

To assess our ability to predict phenotypic evolution from genomic data, we calculated “offsets” for both kinds of data, describing the level of genetic and phenotypic maladaptation relative to a well-adapted reference population [52, 53]. Offsets were calculated for each geographic origin and evolution regime separately. Thus, for each background-regime combination, the reference was chosen as the line replicate with the highest laboratory fitness at the temperature to which it had been adapting. As for our estimates of ancestral thermal performance (Figure 1), we here defined our proxy of laboratory fitness as the ratio between lifetime adult offspring production and their egg-to-adult development time, roughly corresponding to the line’s intrinsic population growth rate. Thus, for example, for hot-adapted lines, Yemen 8 had higher fitness (3.65) than Yemen 7 (2.88) at 35°C and was chosen as the reference line for all 6 Yemen populations phenotyped at 35°C.

For our fitness offset, we used our metric of laboratory fitness, 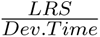, to calculate relative fitness of each tested line relative to its reference line. Phenotypic divergence offsets were calculated as the Euclidean distance in the 7 mean-scaled traits between each tested line and its reference. The set of SNPs used to calculate genomic offsets were those putatively selected for in the reference line (more extreme than the 0.01th or 99.99th quantiles of the null drift distributions, see above). Genomic offsets were calculated as the sum of absolute differences between the reference and tested lines. The difference in each SNP’s allele frequency was scaled by the absolute selection coefficient of that SNP calculated in the reference population such that SNPs with large differences in allele frequency but relatively small selection coefficients would not contribute greatly to the genomic offset.

## Acknowledgements

This work was funded by grants 2019-05024 from the Swedish Research Council (VR) and 2022-01117 from Formas to DB, and Swedish Research Council 2017-04963 and 2022-03427 to RS. The storage and computations for this work were enabled by resources in projects SNIC 2021/5-125 and SNIC 2020/6-175 provided by the Swedish National Infrastructure for Computing (SNIC) at UPPMAX, partially funded by the Swedish Research Council through grant agreement no. 2018-05973. Special thanks to Kalle Tunström and Rachel Steward for figure critiques/suggestions, and Kevin Roberts and Loke von Schmalensee for insight into the physiological limits of thermal adaptation.

## Data and Code Availability Statements

Data and code for analyses are available at Dryad (doi:10.5061/dryad.bzkh189kd) to be released upon acceptance. Accession numbers will be provided upon acceptance for publication. All scripts and data fall under a CC0 1.0 Universal (CC0 1.0) license. Peer-review link for code and data accessibility are available here: http://datadryad.org/stash/share/ioNUncvnoFT2hGKqcIYFKKIunBITTfC75 W0HMzjmFCo

## Supplemental Material, Figures, and Tables

**Table S1:**
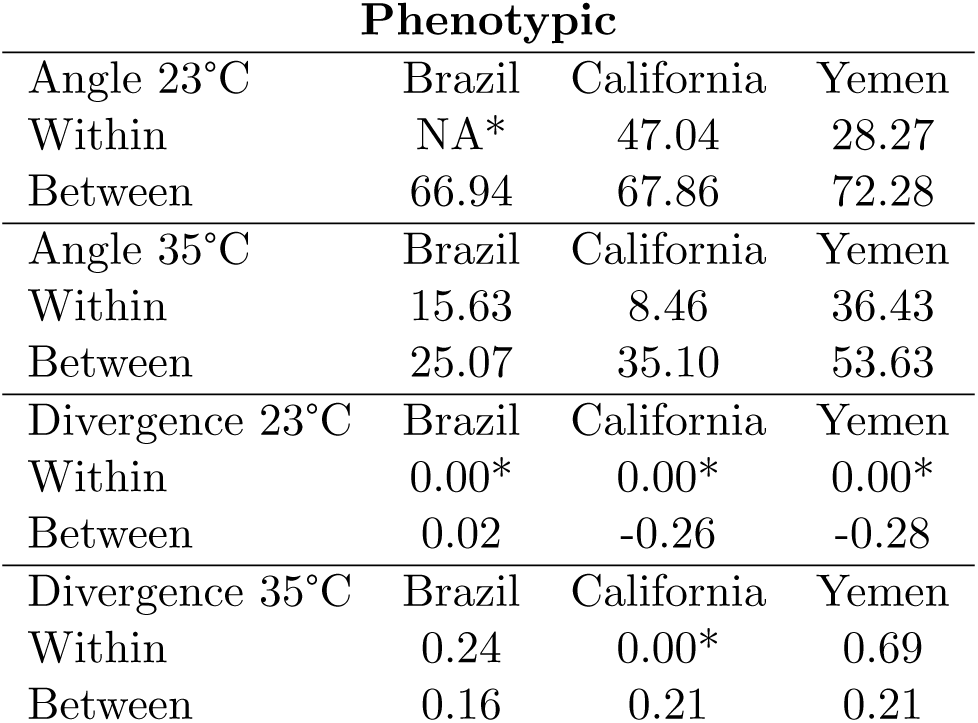
Mean phenotypic angles (*θ*) and divergence (*E_d_ S_d_*) between lines of the same (within) and different (between) genetic background, for each selection regime. Within-background comparisons are represented by a single value (between the two replicates per origin). Asterisks represent estimates with high measurement error relative to vector length.

| **Phenotypic** |  |  |  |
| --- | --- | --- | --- |
| Angle 23°C | Brazil | California | Yemen |
| Within | NA* | 47.04 | 28.27 |
| Between | 66.94 | 67.86 | 72.28 |
| Angle 35°C | Brazil | California | Yemen |
| Within | 15.63 | 8.46 | 36.43 |
| Between | 25.07 | 35.10 | 53.63 |
| Divergence 23°C | Brazil | California | Yemen |
| Within | 0.00* | 0.00* | 0.00* |
| Between | 0.02 | -0.26 | -0.28 |
| Divergence 35°C | Brazil | California | Yemen |
| Within | 0.24 | 0.00* | 0.69 |
| Between | 0.16 | 0.21 | 0.21 |

**Table S2:** Estimated *Ne*, number of putatively selected SNPs, number of putatively selected genes (genes with SNPs falling within protein coding regions), and the mean magnitude of allele frequency change, allele frequency change per generation, and selection coefficient per population. The magnitudes of allele frequency change and selection coefficient estimates are comprised of the putatively selected SNPs identified for each respective line.

| | $Ne$ | # Putative Sites | # Putative Genes | $\vert \Delta AF\vert $ | $\frac{\vert \Delta AF\vert }{gen}$ | $\vert S\vert $ |
| --- | --- | --- | --- | --- | --- | --- |
| Yemen1 (23°C) | 244.29 | 61,458 | 2,405 | 0.546 | 0.009 | 0.0515 |
| Yemen2 (23°C) | 291.21 | 56,378 | 2,170 | 0.511 | 0.008 | 0.0490 |
| Cali3 (23°C) | 291.33 | 86,561 | 2,805 | 0.484 | 0.008 | 0.0440 |
| Cali4 (23°C) | 316.67 | 60,384 | 2,481 | 0.501 | 0.008 | 0.0374 |
| Brazil5 (23°C) | 194.23 | 34,216 | 1,490 | 0.607 | 0.010 | 0.0495 |
| Brazil6 (23°C) | 215.69 | 41,629 | 1,784 | 0.592 | 0.010 | 0.0448 |
| Yemen7 (35°C) | 178.38 | 32,804 | 1,343 | 0.663 | 0.010 | 0.0540 |
| Yemen8 (35°C) | 106.44 | 21,716 | 774 | 0.826 | 0.014 | 0.0660 |
| Cali9 (35°C) | 189.95 | 42,695 | 1,717 | 0.618 | 0.010 | 0.0552 |
| Cali10 (35°C) | 193.84 | 43,445 | 1,731 | 0.610 | 0.010 | 0.0480 |
| Brazil11 (35°C) | 161.86 | 28,758 | 1,217 | 0.658 | 0.010 | 0.0602 |
| Brazil12 (35°C) | 184.51 | 21,551 | 1,146 | 0.706 | 0.011 | 0.0596 |

**Table S3:** Estimated Pearson’s correlation (*r*), between allele frequency change and estimated selection coefficient.

| | $r$ |
| --- | --- |
| Yemen1 (23°C) | 0.47 |
| Yemen2 (23°C) | 0.46 |
| Cali.3 (23°C) | 0.47 |
| Cali.4 (23°C) | 0.49 |
| Brazil5 (23°C) | 0.54 |
| Brazil6 (23°C) | 0.54 |
| Yemen7 (35°C) | 0.47 |
| Yemen8 (35°C) | 0.59 |
| Cali.9 (35°C) | 0.48 |
| Cali.10 (35°C) | 0.50 |
| Brazil11 (35°C) | 0.53 |
| Brazil12 (35°C) | 0.52 |

**Table S4:**
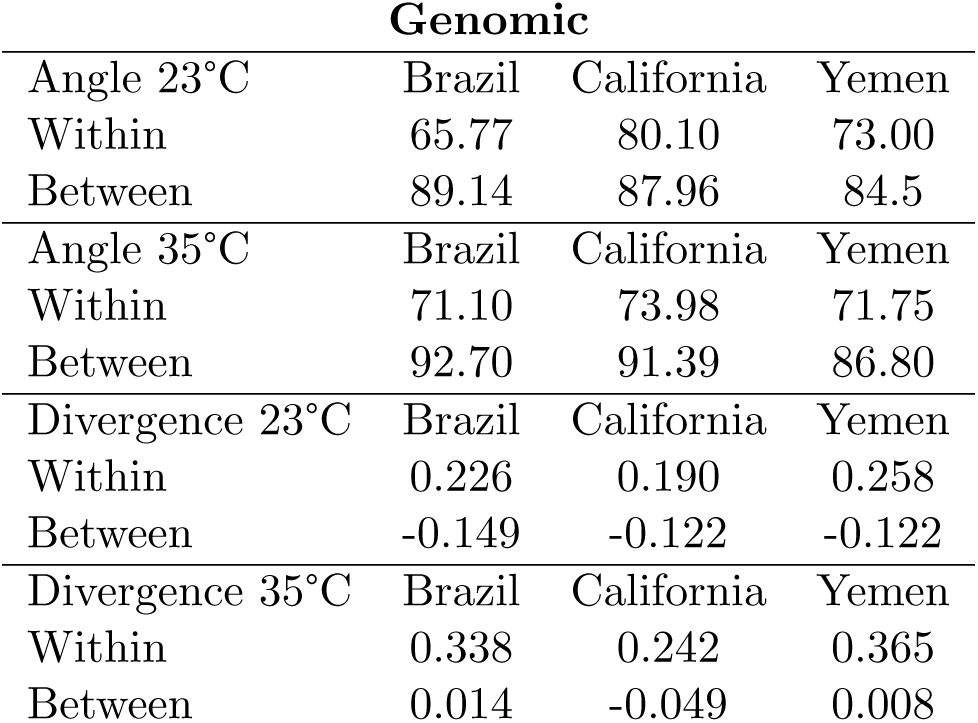
Mean genomic angles (*θ*) and divergence (*E_d_ S_d_*) between lines of the same (within) and different (between) genetic background, for each selection regime. Within-background comparisons are represented by a single value (between the two replicates per origin).

**Table S5:** Gene set enrichment analysis for genic targets of selection shared across all 6 cold-adapted replicates (top), and all 6 hot-adapted replicates (bottom).

| GO.ID 23°C | Term | Ann. | Sig. | Exp. | Adj. p-value |
| --- | --- | --- | --- | --- | --- |
| GO:0035556 | intracellular signal... | 195 | 13 | 6.73 | 0.0035 |
| GO:0007156 | homophilic cell adhe... | 22 | 4 | 0.76 | 0.0062 |
| GO:0035023 | regulation of Rho pr... | 29 | 4 | 1.00 | 0.0166 |
| GO:0007205 | protein kinase C-act... | 8 | 2 | 0.28 | 0.0289 |
| GO:0032940 | secretion by cell | 12 | 2 | 0.41 | 0.0344 |
| GO:0007165 | signal transduction | 473 | 32 | 16.34 | 0.0410 |
| GO:0007186 | G protein-coupled re... | 148 | 11 | 5.11 | 0.0487 |
| GO:0040007 | growth | 20 | 8 | 0.69 | 1.5e-07 |
| GO.ID 35°C | Term | Ann. | Sig. | Exp. | Adj. p-value |
| GO:0007156 | homophilic cell adhe... | 22 | 2 | 0.12 | 0.006 |
| GO:0007018 | microtubule-based mo... | 46 | 2 | 0.25 | 0.025 |
| GO:0019319 | hexose biosynthetic ... | 5 | 1 | 0.03 | 0.027 |
| GO:0006013 | mannose metabolic pr... | 5 | 1 | 0.03 | 0.027 |

**Table S6:** Mean genomic angles (*θ*) and divergence (*E_d_ S_d_*) of within- and betweenbackground comparisons for each selection regime using selected sites which were variable in the ancestors of all populations. Within-comparisons are represented by a single value between the two replicates per origin.

| **Shared Genomic** |  |  |  |
| --- | --- | --- | --- |
| Angle 23°C | Yemen | California | Brazil |
| Within | 74.56 | 75.29 | 71.67 |
| Between | 81.87 | 83.45 | 84.03 |
| Angle 35°C | Yemen | California | Brazil |
| Within | 70.52 | 75.07 | 69.19 |
| Between | 84.69 | 89.15 | 90.49 |
| Divergence 23°C | Yemen | California | Brazil |
| Within | 45.53 | 34.07 | 36.98 |
| Between | 3.9 | 8.25 | 7.6 |
| Divergence 35°C | Yemen | California | Brazil |
| Within | 60.36 | 44.34 | 57.08 |
| Between | 22.39 | 17.12 | 18.63 |

**Figure S1:**
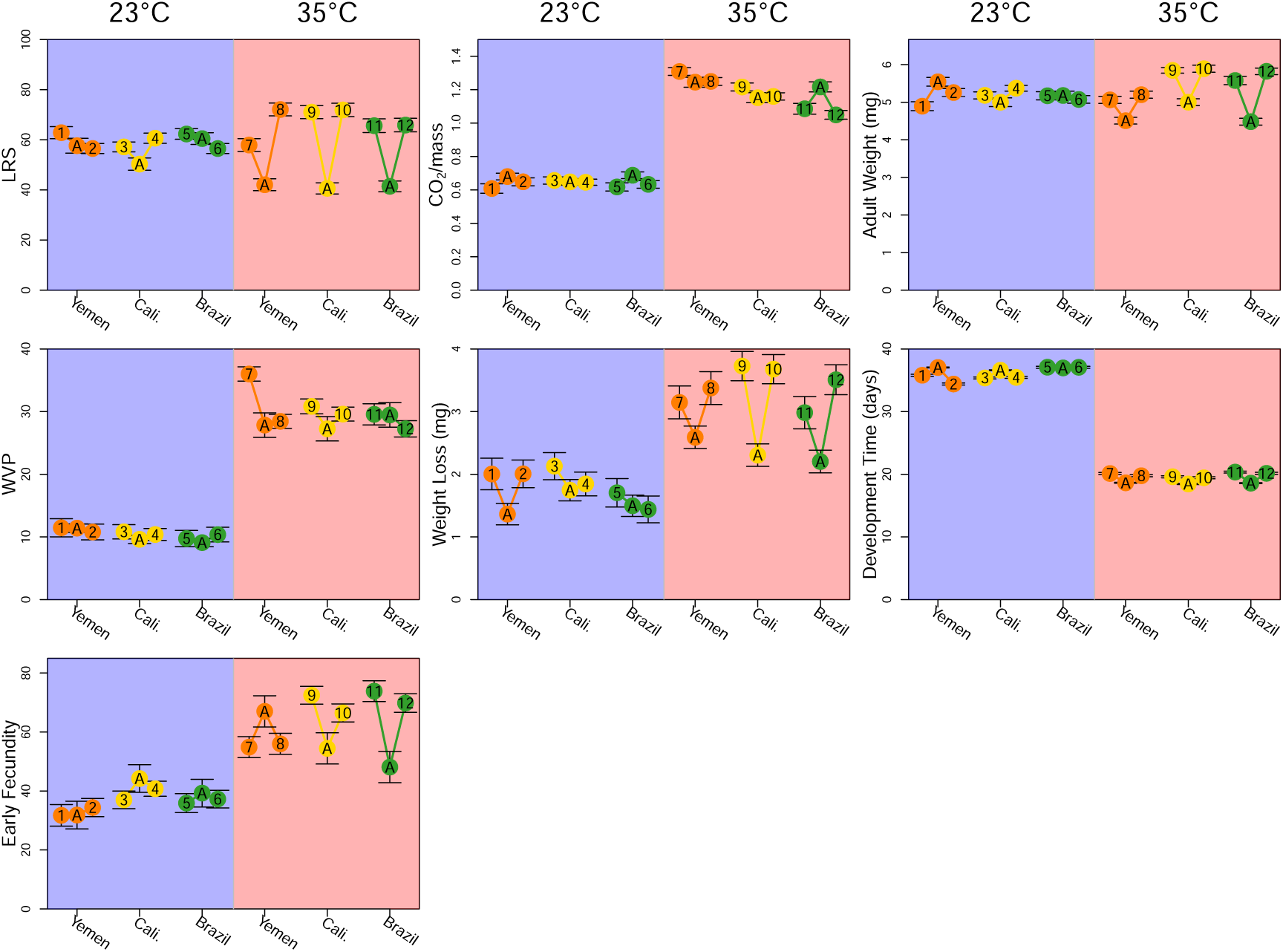
Means and standard errors for the seven life-history traits in evolved and ancestral lines. Note that ancestors (”A”) and evolved lines (1–12) were measured in their evolved temperature regime, denoted by background color.

**Figure S2:**
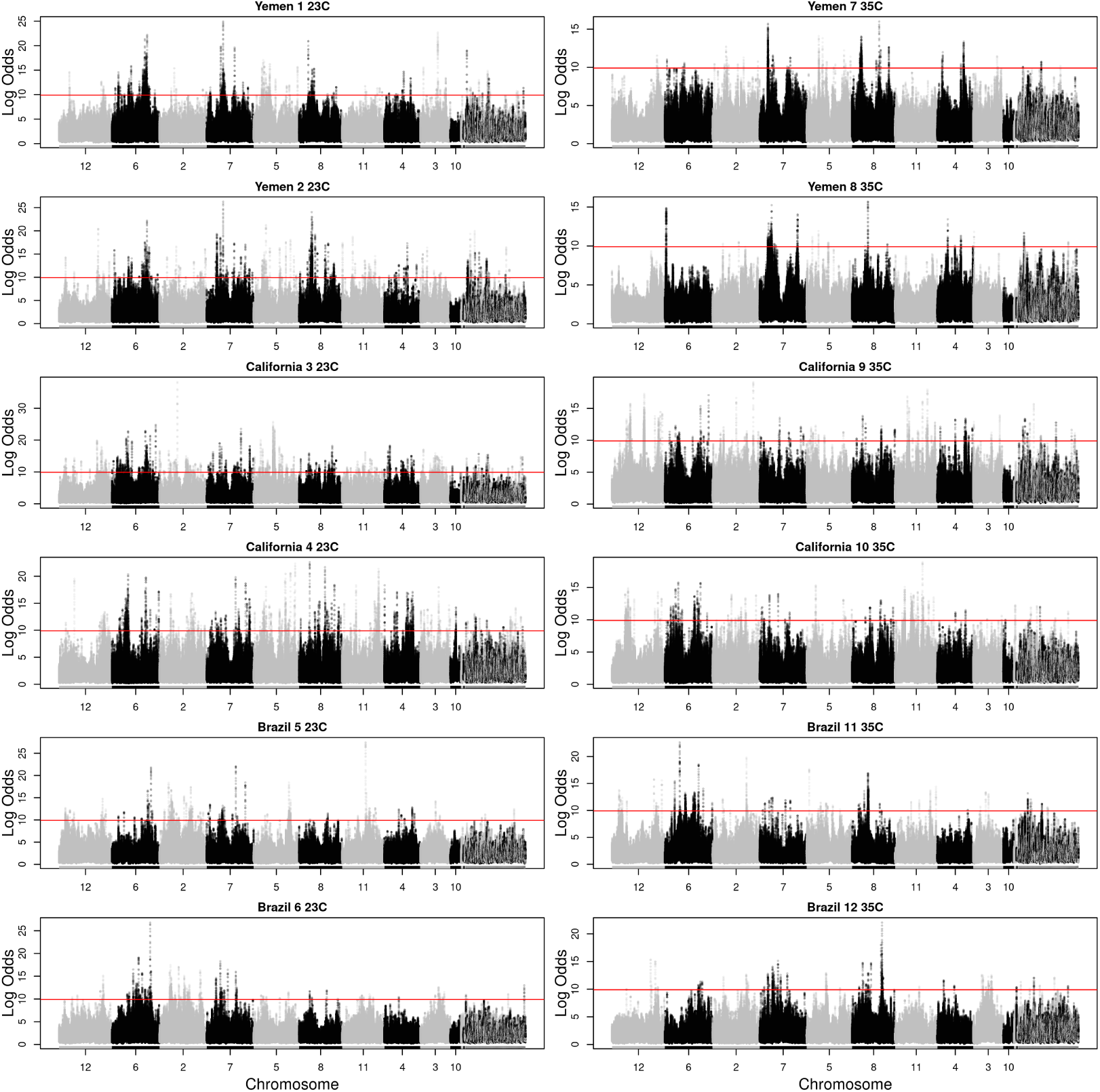
Manhattan plots of rolling average log odds p-values (window size = 20 SNPs) for each line across all selected sites using R package RcppRoll (v.0.3.0). The largest 10 scaffolds (i.e., chromosomes) are named. The significance threshold of log odds (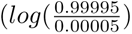) is given by the horizontal red line.

**Figure S3:**
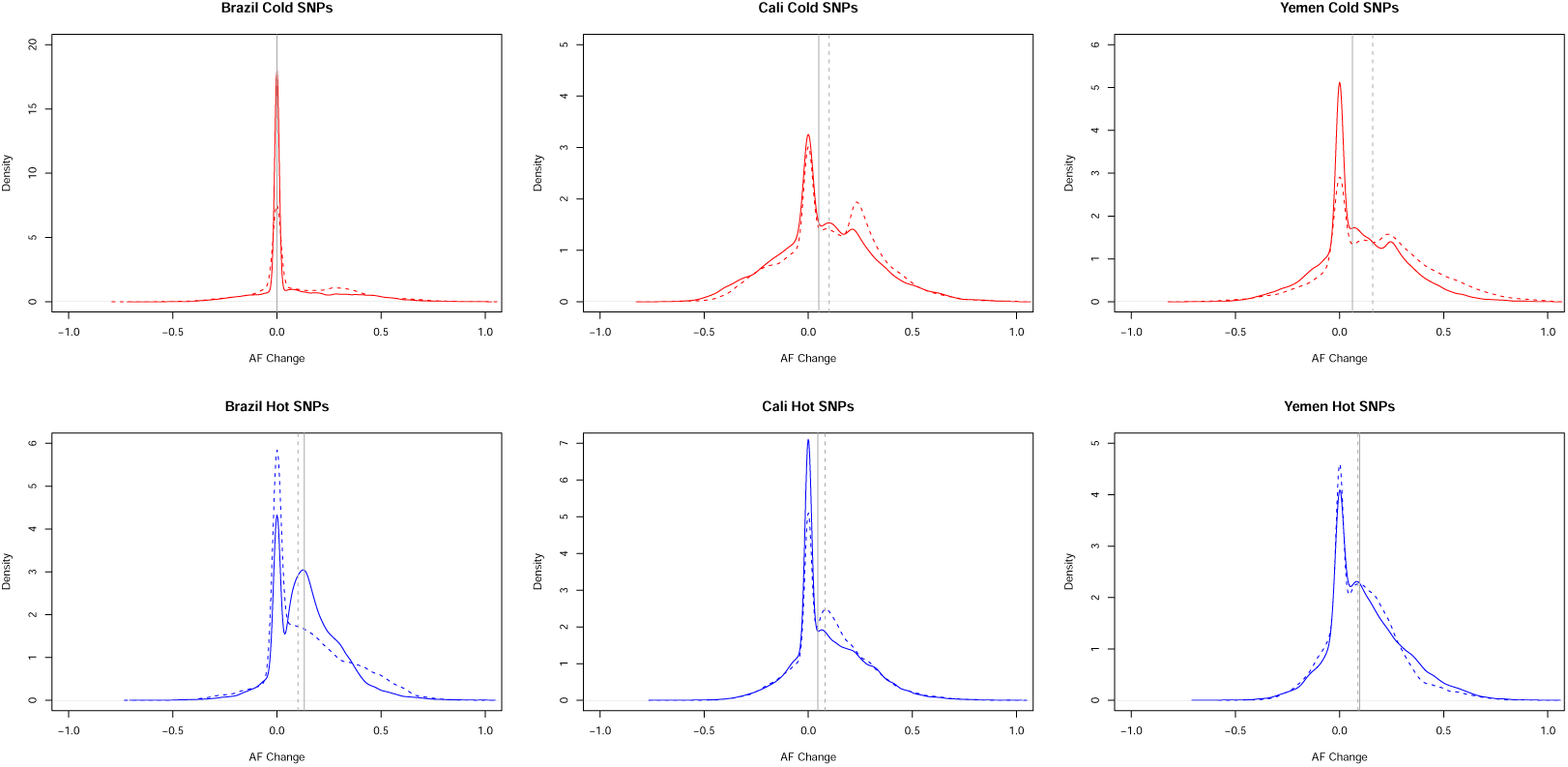
Allele frequency changes in hot (red) and cold (blue) lines for candidate SNPs identified in the alternative thermal regime. For each selected SNP in a thermal regime, we quantified allele frequency changes in the alternative thermal regime (i.e., cold-associated SNPs in heat-adapted lines, and hot-associated SNPs in cold-adapted lines). Allele frequency changes were calculated such that positive values indicate a change in the same direction as found in the thermal regime where the SNP was identified as under putative selection (implying synergistic effects on fitness across temperatures), whereas negative values indicate a change in the opposite direction (implying antagonistic effects on fitness across temperatures). The two replicate lines per background are shown separately as solid and dashed colored lines. Vertical solid and dashed grey lines indicate the median allele frequency change of the respective replicate line. While there are many examples of allele frequency changes going in both the same and opposite direction in the two thermal regimes, there is a large over-representation of selected SNPs showing no change in the alternative thermal regime, and the mean change even seems to be slightly positive. This suggests that SNPs involved in adaptation during the experiment tended to have effects that were private to each temperature and provides little evidence for a dominating role of antagonistic pleiotropy

**Figure S4:**
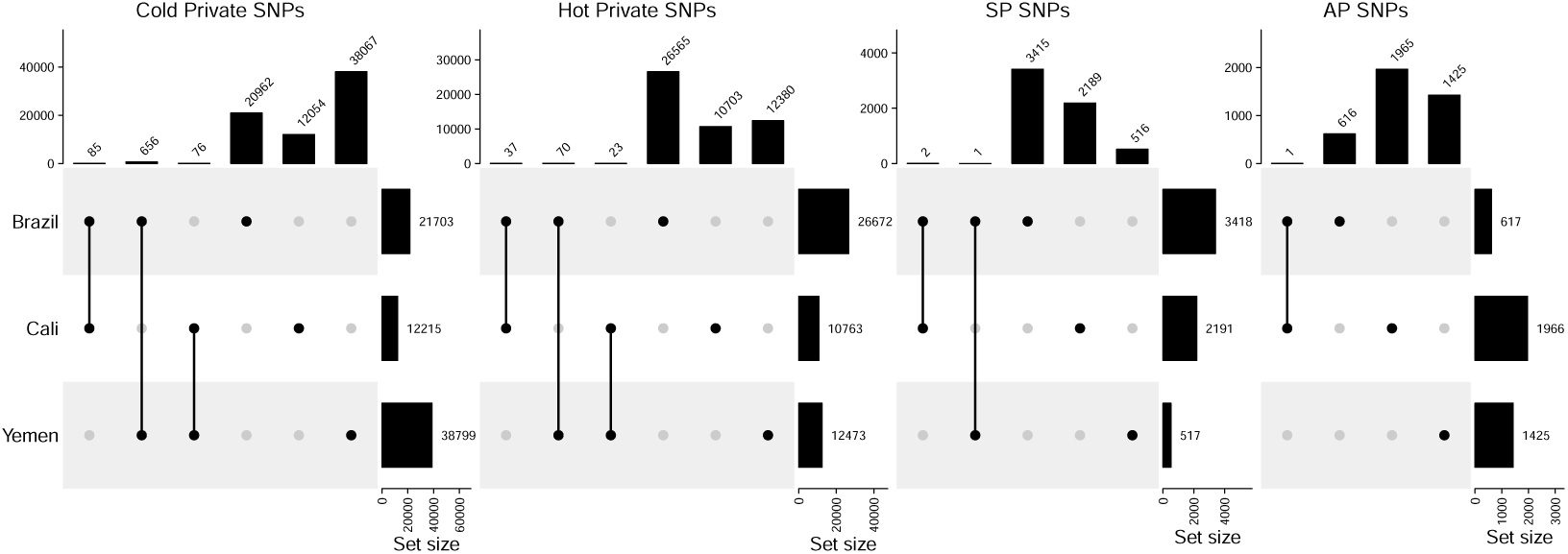
Upset plots of the different categories of SNPs; Cold Private (selected only at cold), Hot Private (selected only at hot), AP (antagonistically pleiotropic - selected in opposite directions in the hot and cold regime) and SP (Synergistically pleiotropic - selected in the same direction in the hot and cold regime).

**Figure S5:**
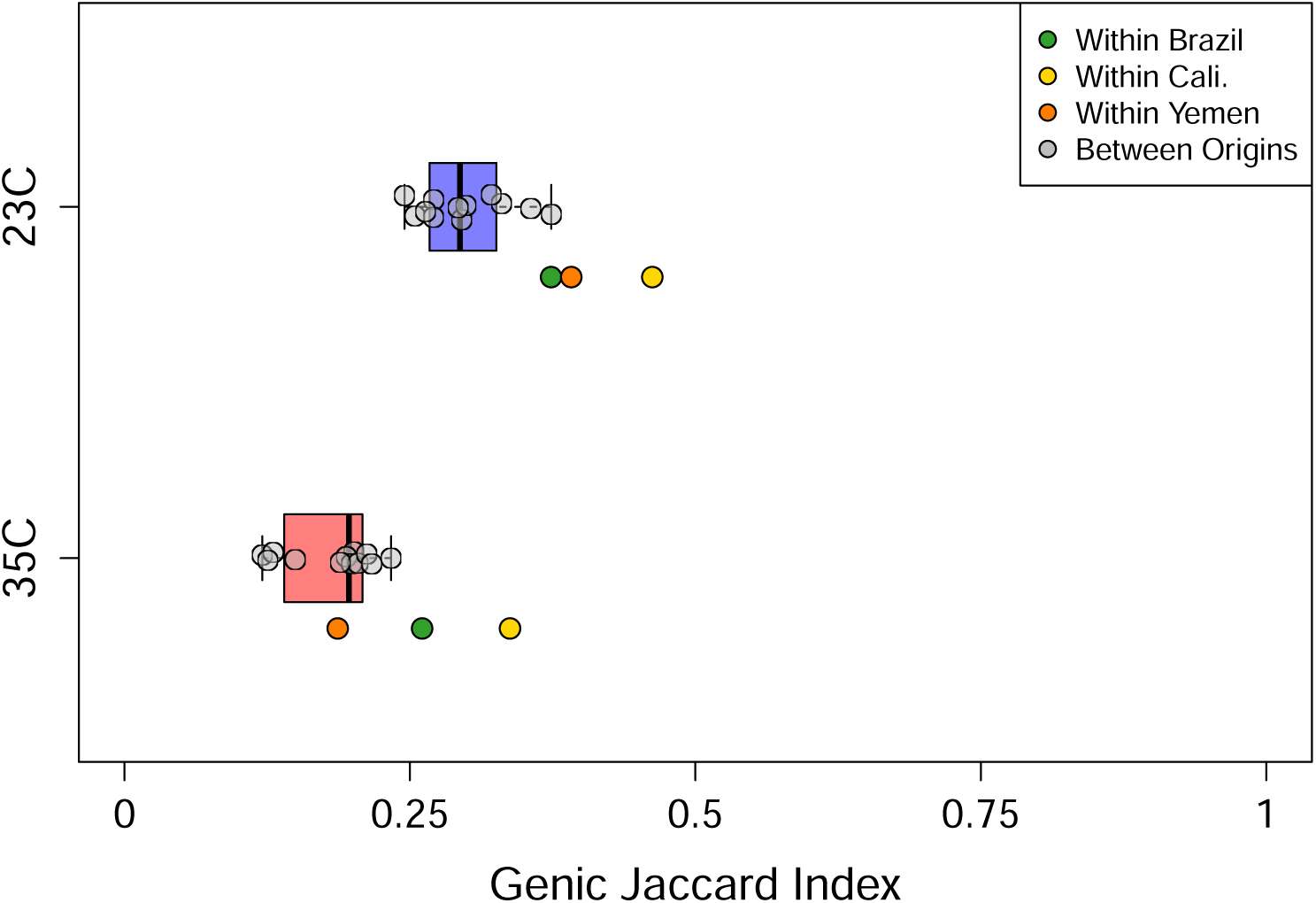
Distribution of Jaccard indices for genic targets of selections between lines within temperature regimes. Distributions of angles and divergences are separated by between-origin (grey points) within-origin (colored points) pairwise comparisons. A Jaccard index of 0 indicates no overlap in genic targets, while an index of 1 indicates complete overlap in genic targets.

**Figure S6:**
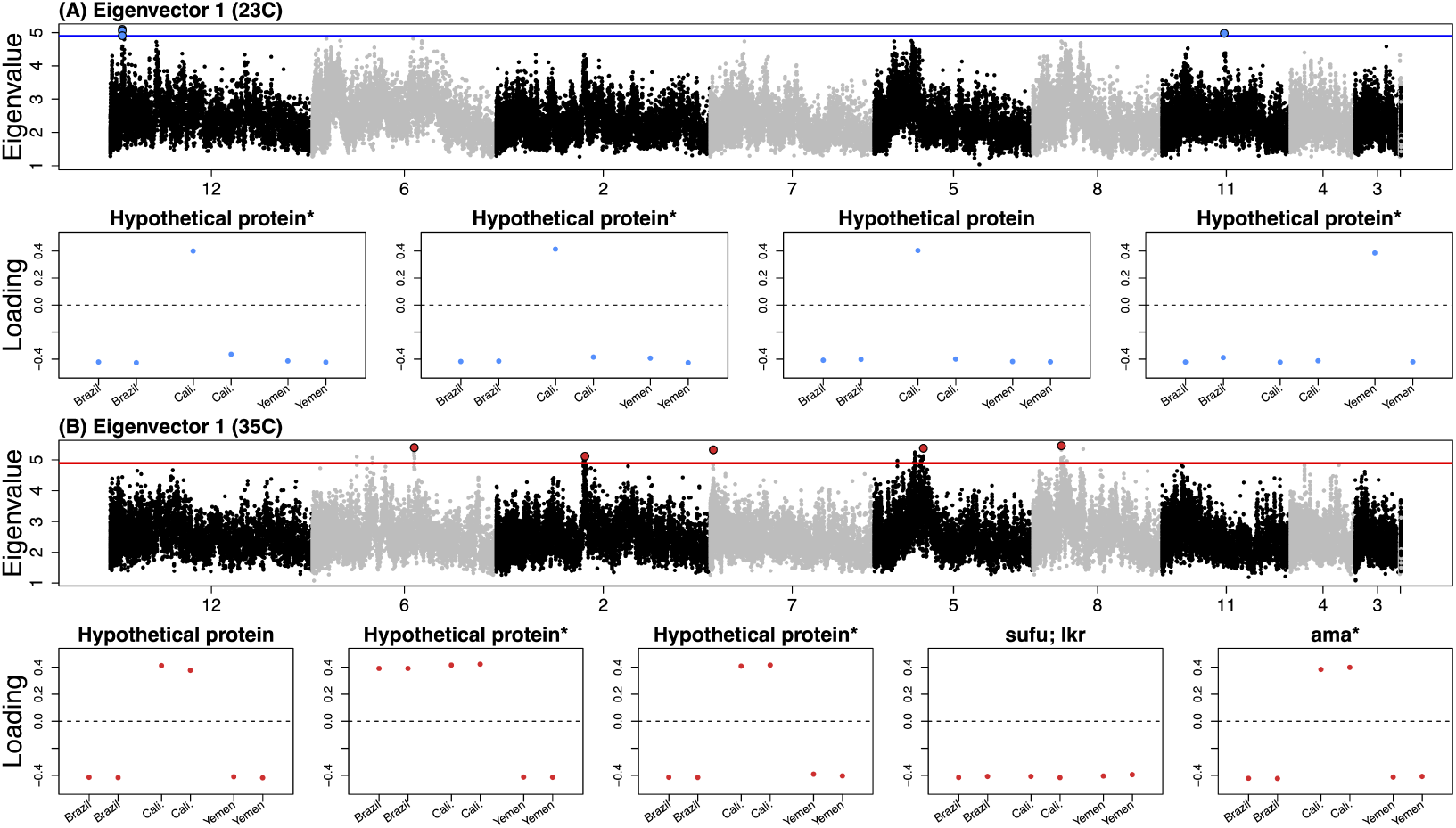
Eigen analyses of genomic regions using 200-SNP windows per regime. Windows exceeding the 99% significance threshold are highlighted– the top 5 independent chromosomal windows for 35°C and all 4 significant windows for 23 °C. For each significant window (presented in left-to-right order in the genome), we show population loadings on Eigenvector 1, with genes in or nearest to each window labeled (nearest genes marked with asterisks).

**Figure S7:**
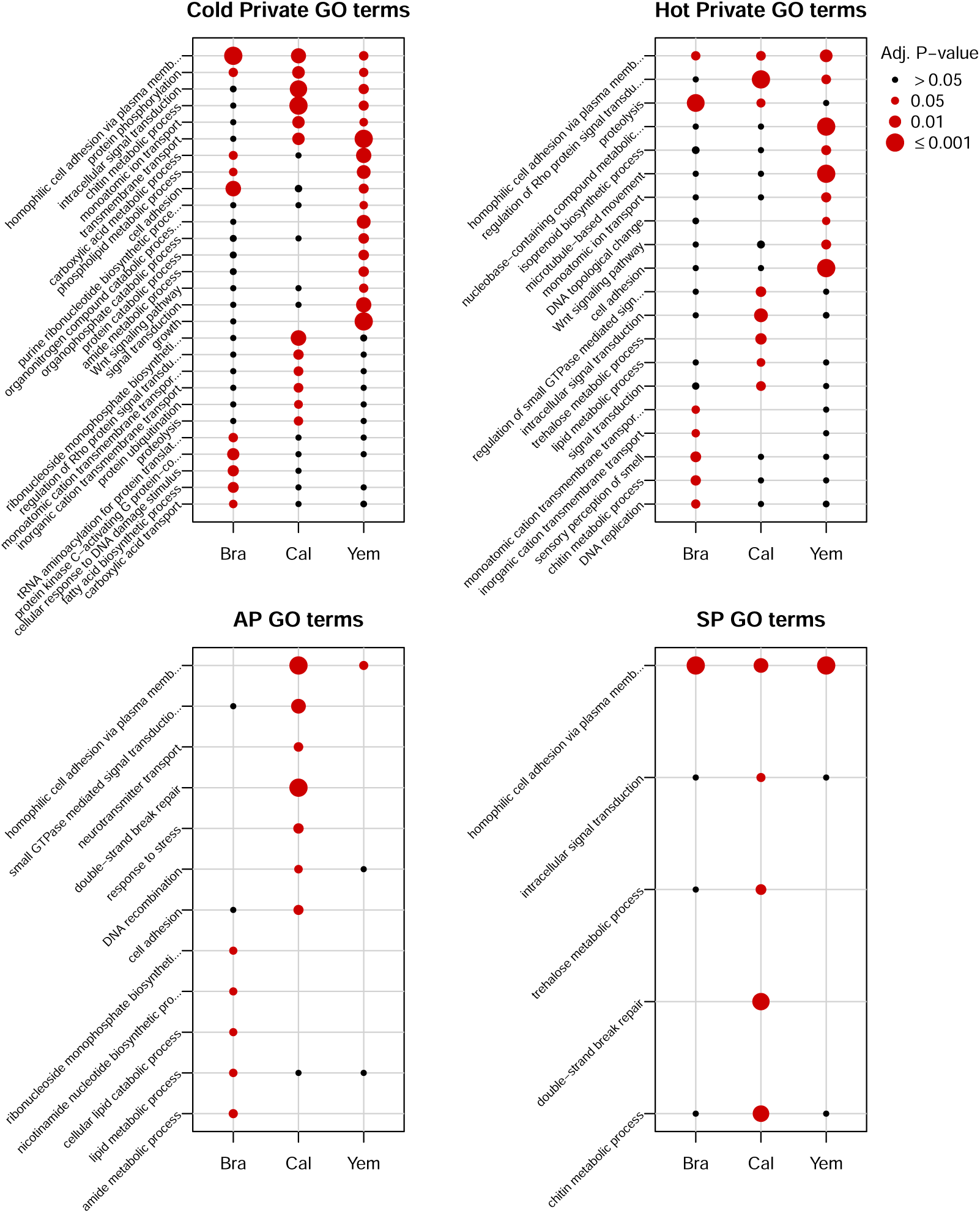
Dot plot indicating significant GO terms per gene set category (based on Fig. 4). Adjusted p-values equal to 1 are not shown.

**Figure S8:**
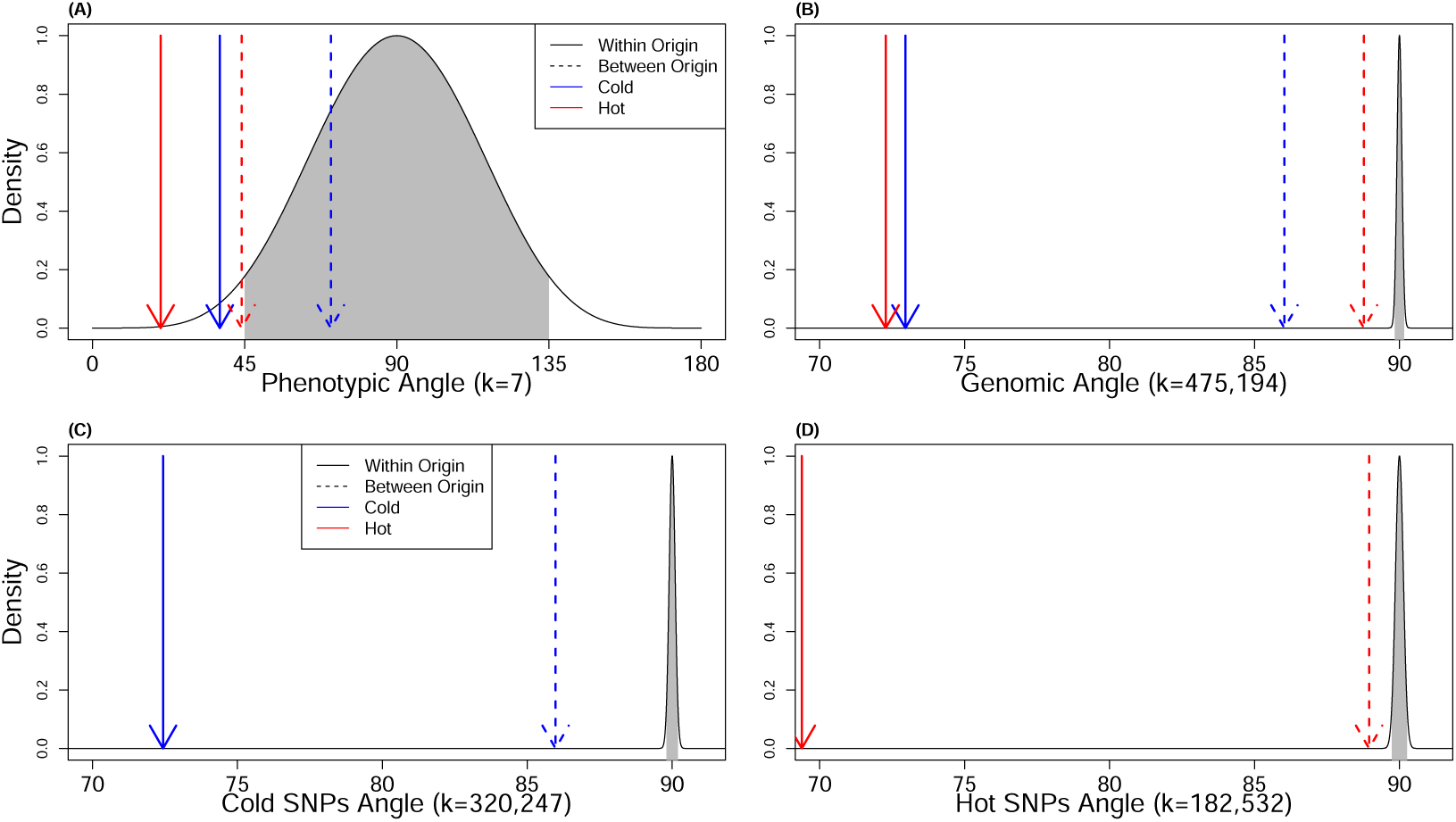
Null distributions of random angles in k-dimensional space associated with (A) phenotypic parallelism, (B) genomic parallelism using any selected site, and (C & D) genomic parallelism using sites selected for per regime. The 95% confidence intervals associated with random angles are shown shaded in grey. The mean within- and between-origin comparisons are presented as arrows and colored per temperature regime.

**Figure S9:**
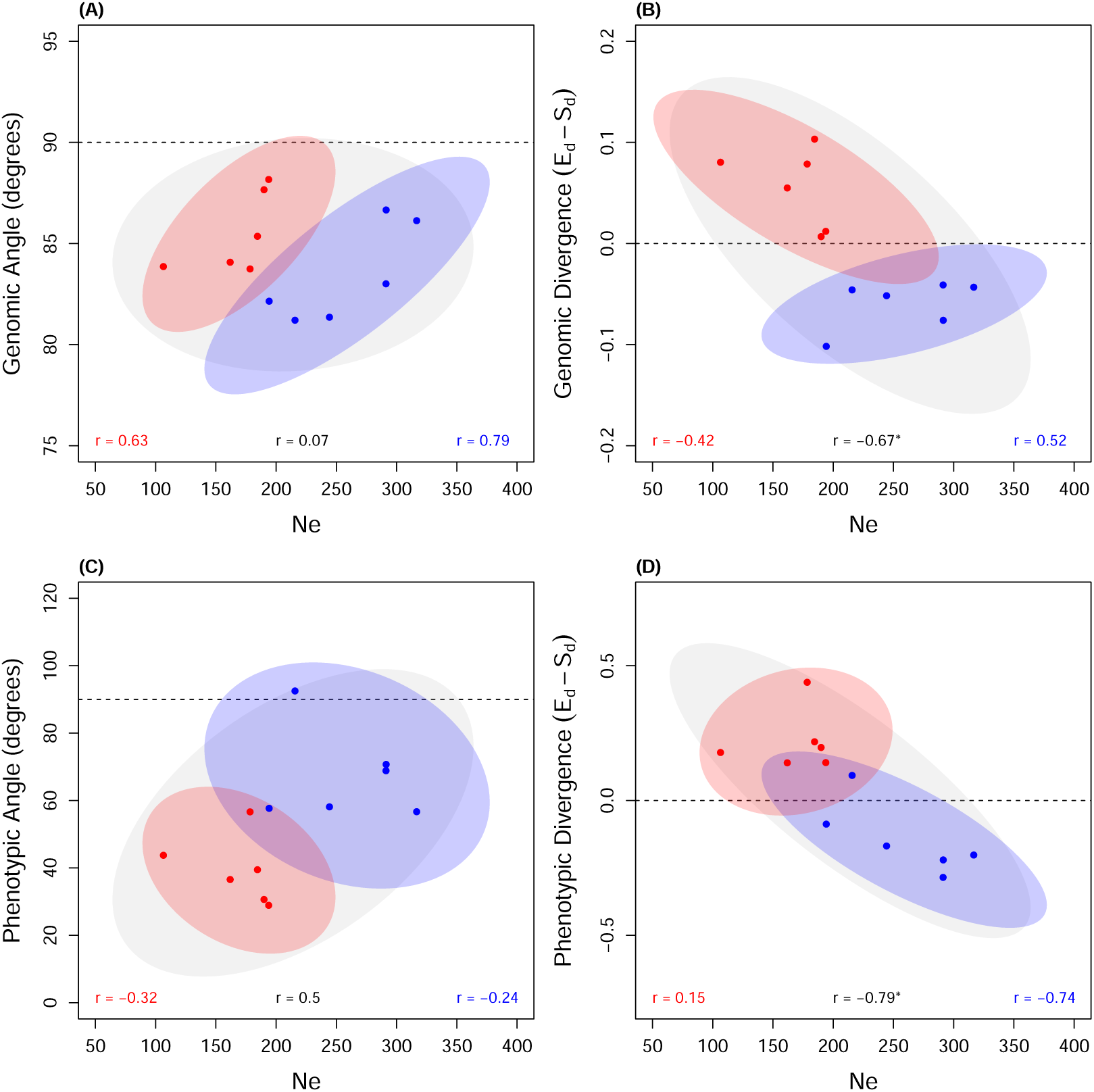
Correspondence between mean measures of phenotypic (A–B) and genomic (C–D) repeatability, and estimates of effective population size (*Ne*) per line. In each plot, points are colored by regime. Confidence ellipses (95%) and correlations are given both per regime (red, 35°C; blue, 23°C) and across regimes (grey) based on line means. Significant correlations are designated with an asterisk.

**Figure S10:**
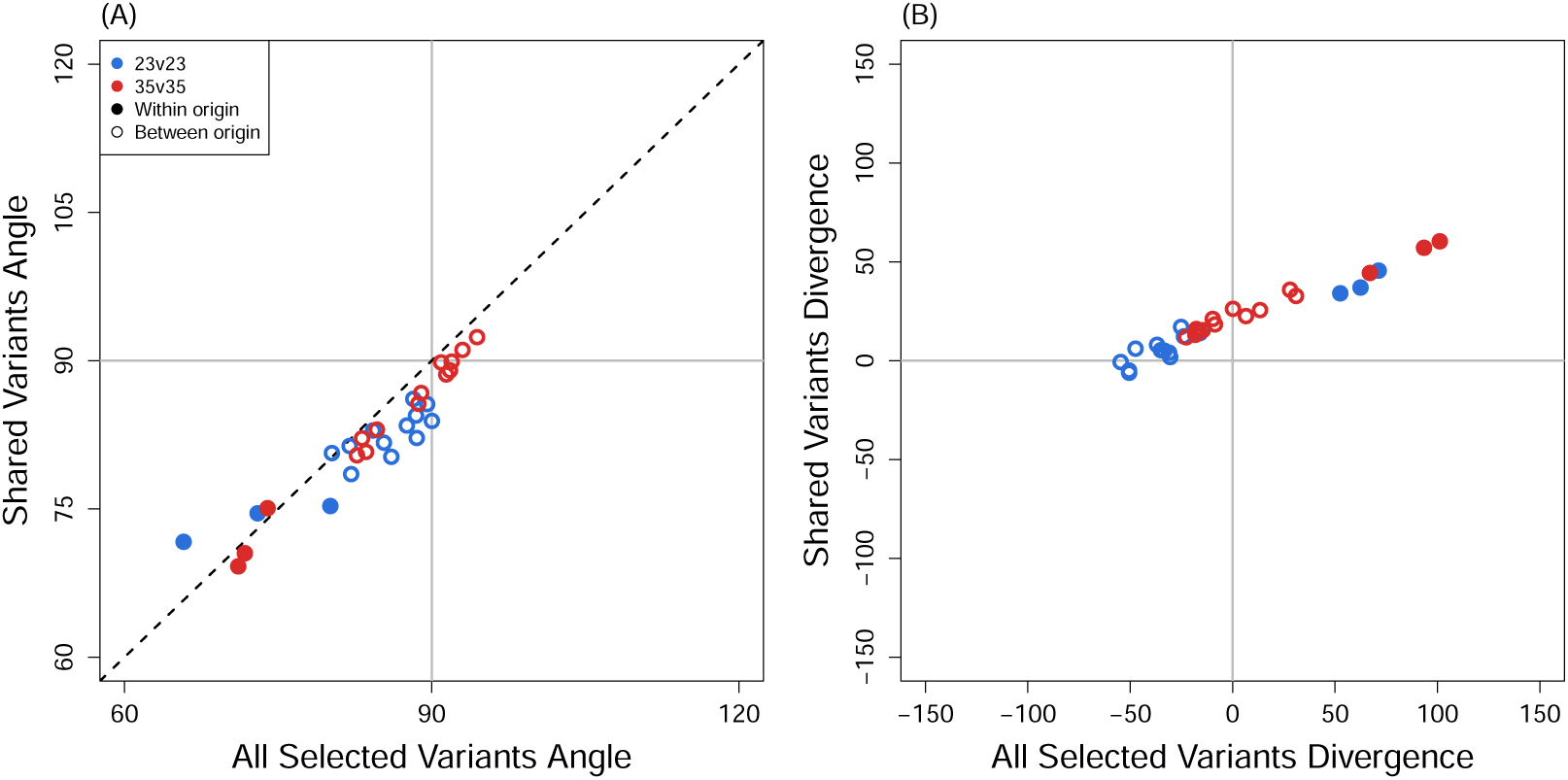
Comparison of repeatability measures between the set of SNPs selected in any one population versus the set of SNPs selected in any one population, but also segregating in all populations. The black dashed line represents the identity line.

**Figure S11:**
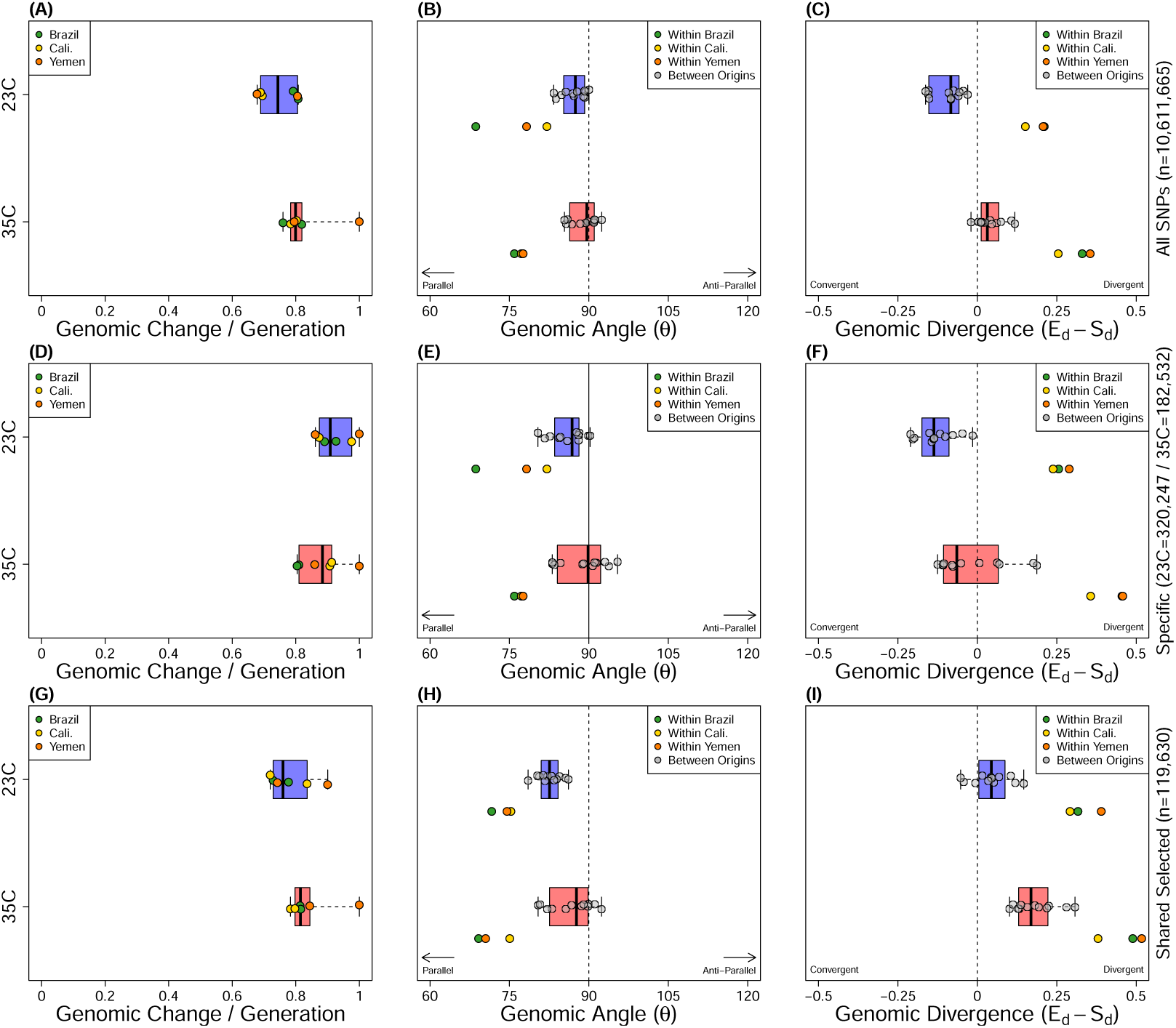
Comparisons between genomic evolutionary rates (A & D & G), pairwise angles (B & E & H) and measurements of convergent or divergent evolution (C & F & I) between different selection criteria for SNP sets. Allele frequency changes found in all SNPs called (top row), SNPs selected for in each temperature regime independently (middle row), and SNPs which are both selected for in any line but also polymorphic among all ancestors (bottom row). Angles and divergences are given for pairwise comparisons between populations of different (grey points) and same (colored points) genetic background. Evolutionary rates have been scaled by the maximum evolutionary rate in the dataset, while divergences have been scaled by the distance between the two most differentiated ancestors.

**Figure S12:**
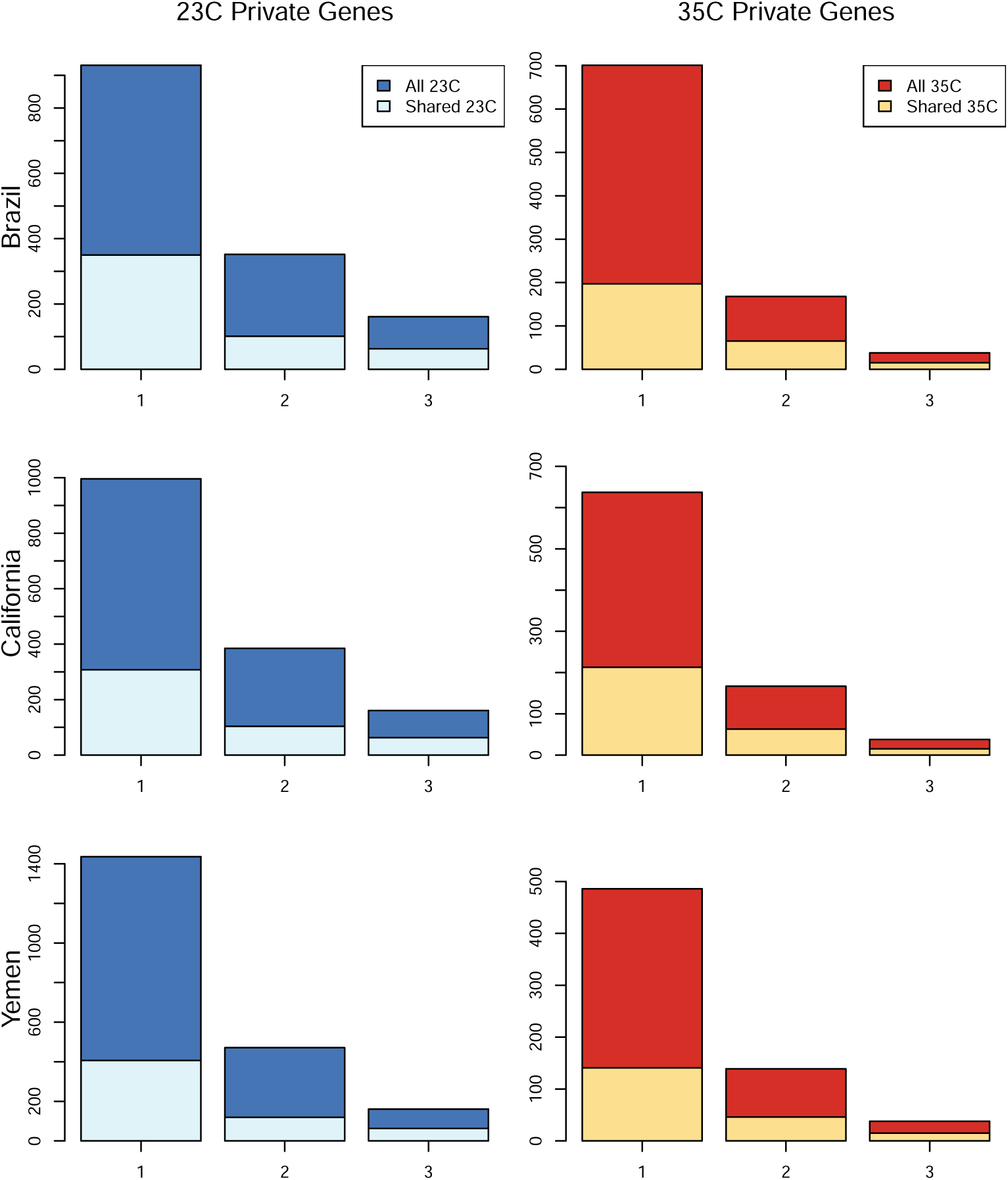
Distributions of the number of privately selected genes shared between populations, for each thermal regime. Genes were found to be under selection in either a single (1) or (2) backgrounds, or on all three backgrounds (3). The distributions for both the entire set of selected sites (n=475,194) and sites which are polymorphic across all ancestral backgrounds (n=119,630) are shown in different shades. Chi-square tests show no significant differences between the distributions for all selected sites versus shared sites for any of the three comparisons (*p >* 0.05).

**Figure S13:**
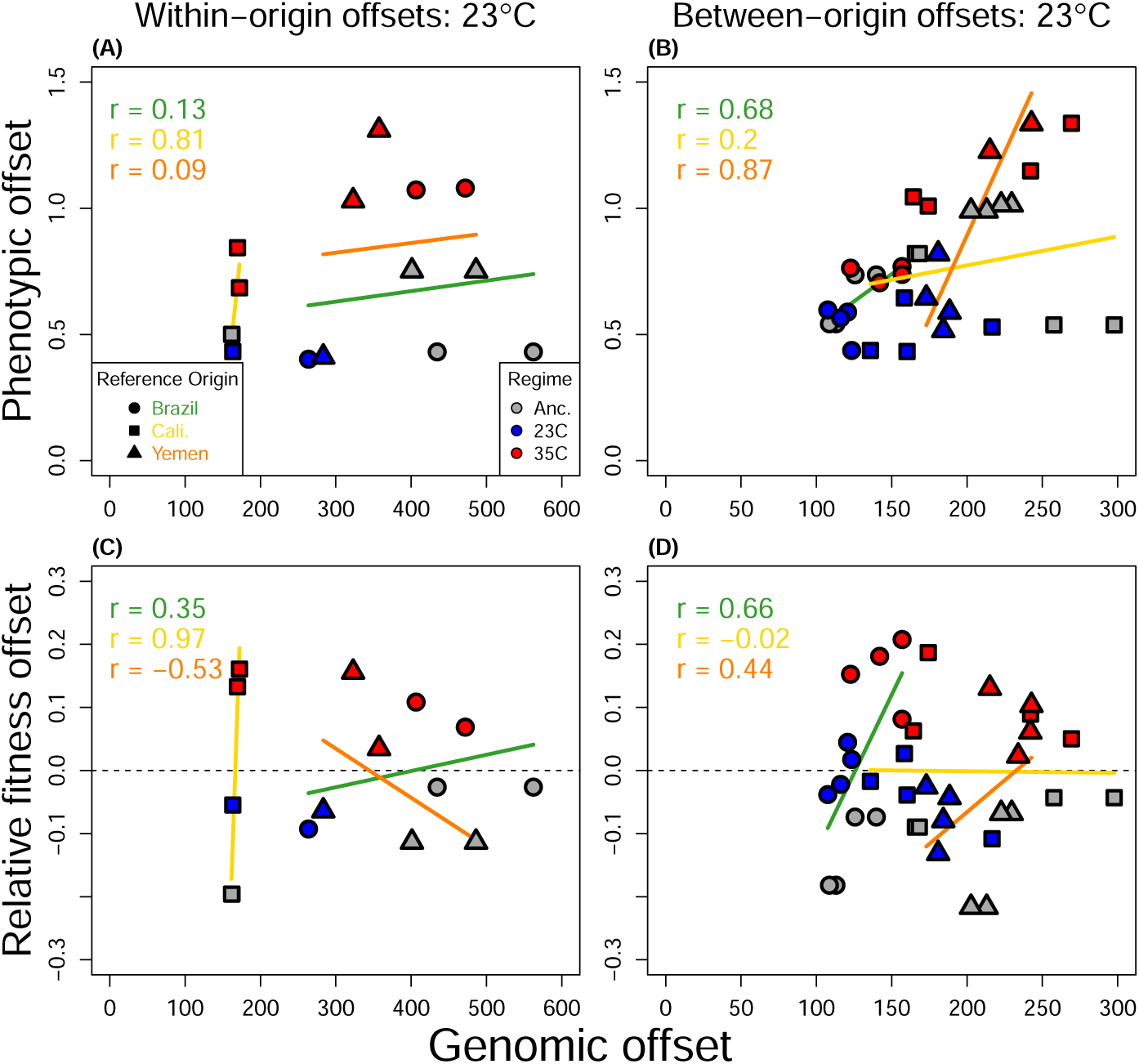
Genomic predictors of phenotypic divergence and relative fitness offsets at cold temperature. Genomic offsets were calculated per reference origin based on SNPs whose p-values for allele frequency change fell within the top 0.001th quantile (n=10546–10887). Phenotypic offsets were calculated as the Euclidean distance in scaled trait-space between the tested line and the reference line. Relative fitness offsets were calculated as the laboratory fitness 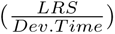 of the tested line relative to that of the reference line. Offsets are organized by within-origin comparisons (A,C) and between-origin comparisons (B,D). Correlations and regression lines are colored by the geographic origin of the reference line.

**Figure S14:**
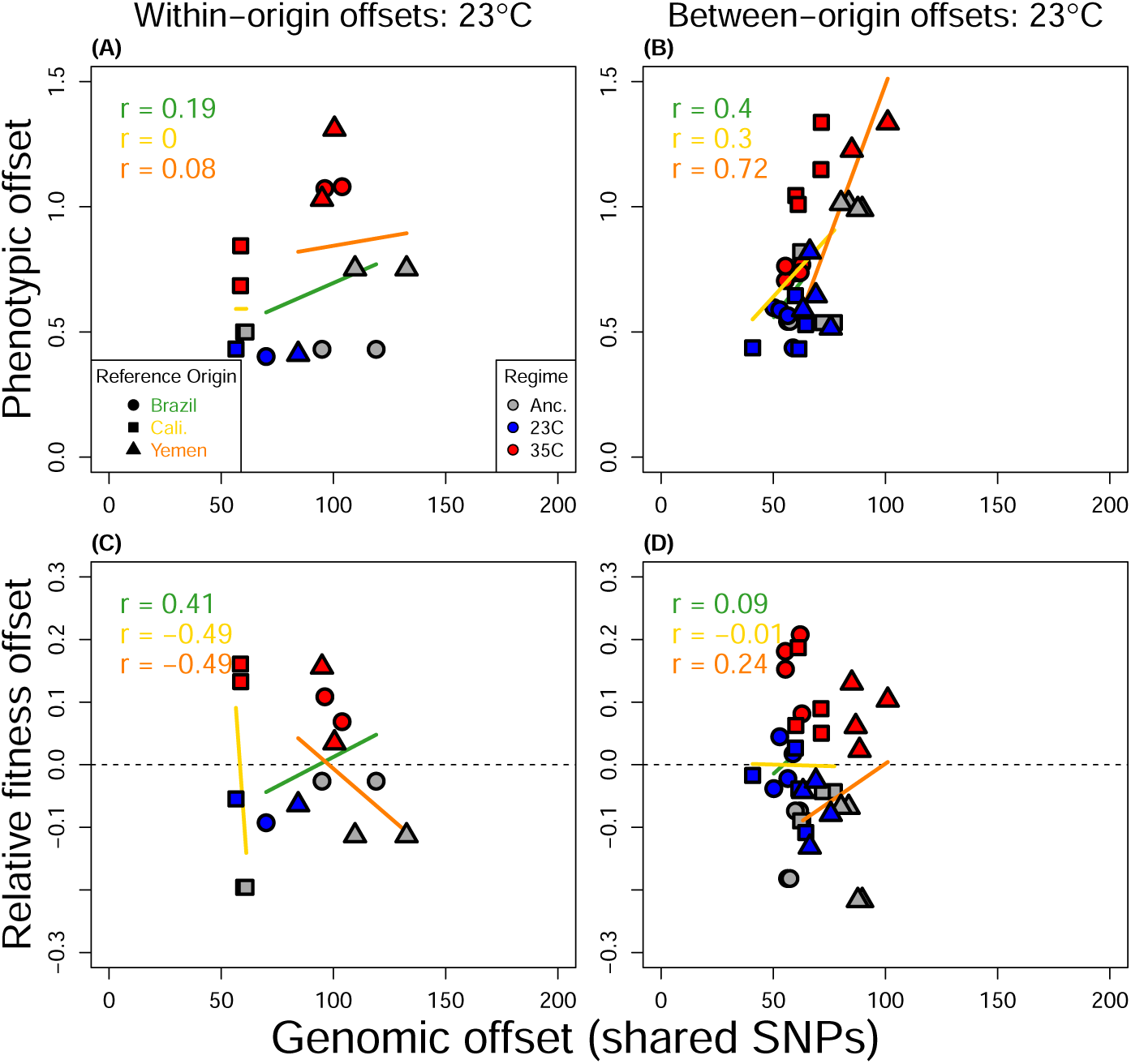
Genomic predictors of phenotypic divergence and relative fitness offsets at cold temperature. Genomic offsets were calculated per reference origin based on SNPs which were polymorphic among all ancestors and whose p-values for allele frequency change fell within the top 0.001th quantile (n=2750–2804). Phenotypic offsets were calculated as the Euclidean distance in scaled trait-space between the tested line and the reference line. Relative fitness offsets were calculated as the laboratory fitness 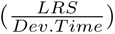 of the tested line relative to that of the reference line. Offsets are organized by within-origin comparisons (A,C) and between-origin comparisons (B,D). Correlations and regression lines are colored by the geographic origin of the reference line.

**Figure S15:**
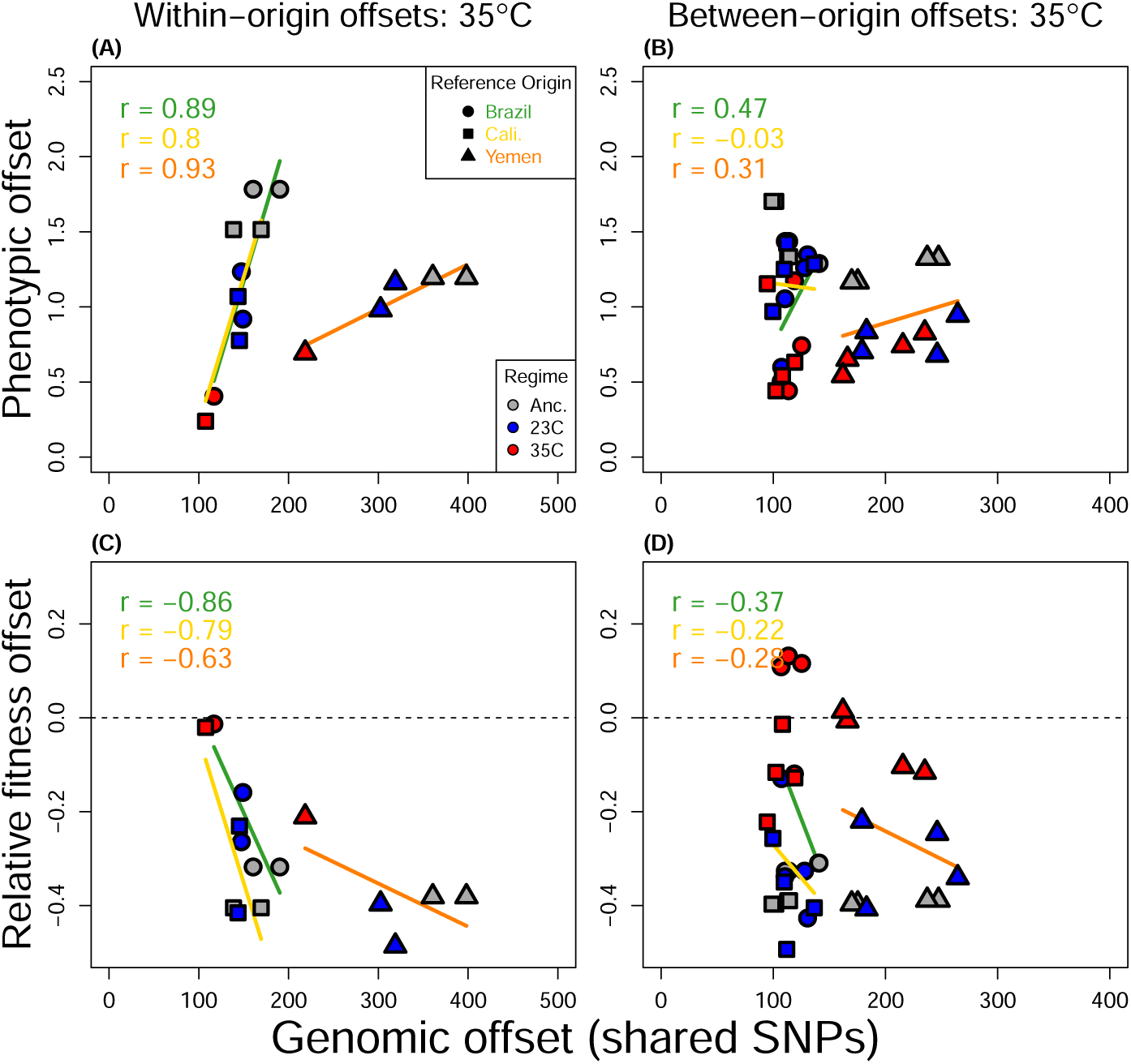
Genomic predictors of phenotypic divergence and relative fitness offsets at hot temperature. Genomic offsets were calculated per reference origin based on SNPs which were polymorphic among all ancestors and whose p-values for allele frequency change fell within the top 0.001th quantile (n=2750–2787). Phenotypic offsets were calculated as the Euclidean distance in scaled trait-space between the tested line and the reference line. Relative fitness offsets were calculated as the laboratory fitness 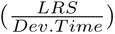 of the tested line relative to that of the reference line. Offsets are organized by within-origin comparisons (A,C) and between-origin comparisons (B,D). Correlations and regression lines are colored by the geographic origin of the reference line.

**Figure S16:**
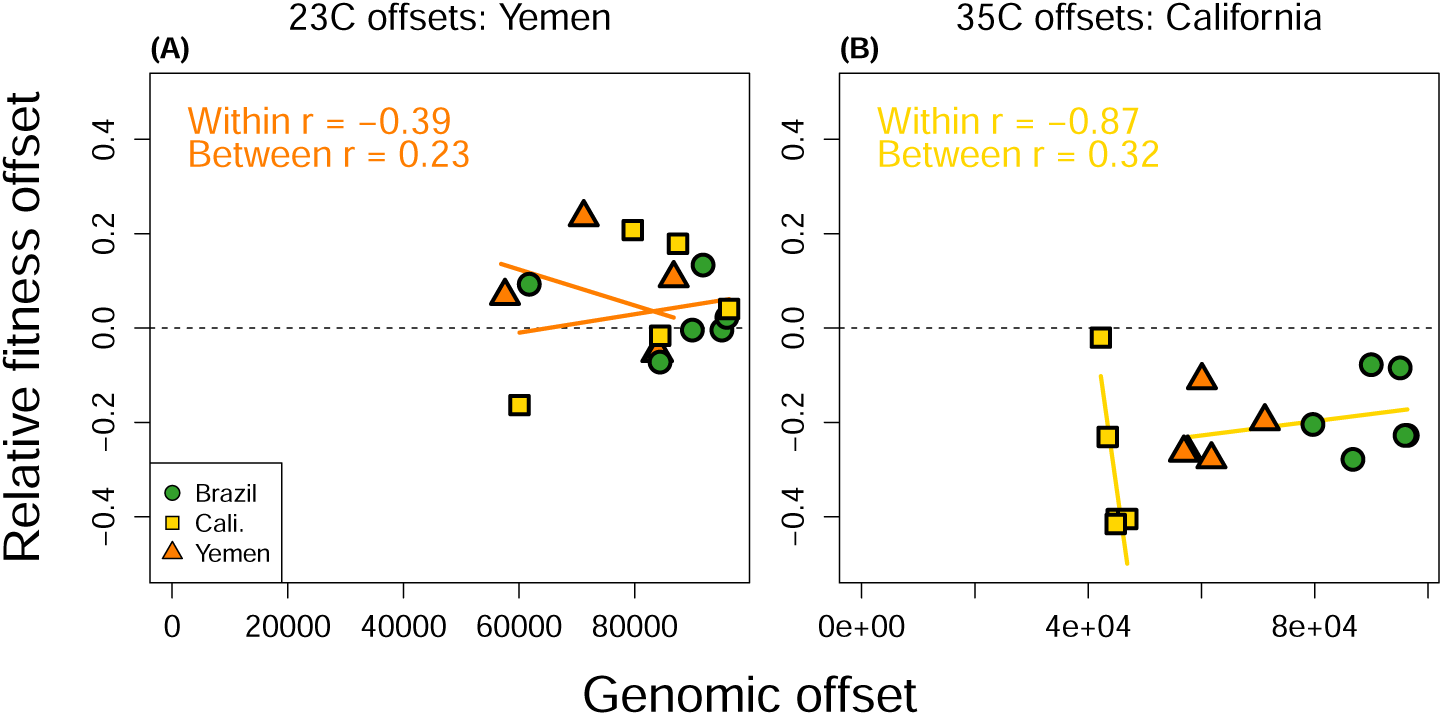
Genomic predictors of relative fitness offsets using lines with the greatest performance per thermal regime as the reference. Genomic offsets were calculated based on all putatively selected SNPs identified in any line replicate, for each thermal regime seperately (Yemen at 23°C, California at 35°C). Allele frequency changes were not scaled by selection coefficients as for other genomic offsets. Relative fitness offsets were calculated as the tested line’s laboratory fitness 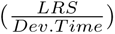 relative to that of the reference line. Correlations and regression slopes are separated by predictions within and between origins.

**Figure S17:**
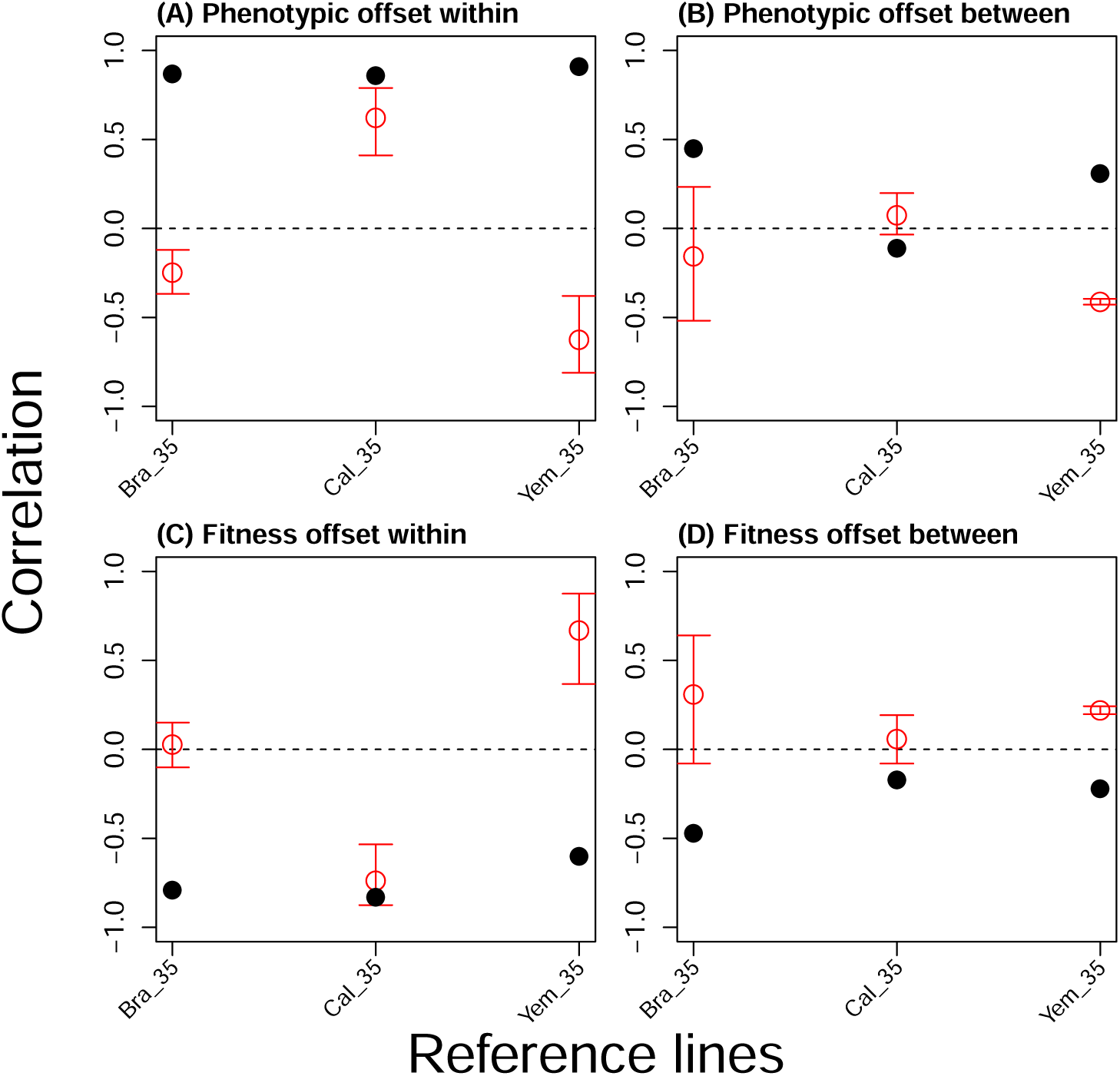
Bootstrapped correlations for genomic offsets at hot temperature using sets of randomly chosen SNPs. Genomic offsets were calculated 1,000 times using randomly sampled SNPs of equal number to the SNPs used in hot line offsets (n=10,546–10,649). The means and 0.025th and 0.975th quantiles for bootstrapped correlations are shown as colored points and lines, respectively. The correlation based on selected SNPs is shown as a solid black point (see Fig. 6. Correlations are separated by within and between origins.

### A hypothesis for how temperature extremes affect epistasis for fitness based on the thermodynamics of enzyme performance

Biological rates of ectotherms show an empirically well-described, roughly exponential, increase with temperature that closely mirrors the thermodynamic properties of enzyme reactions [1, 2]. This pattern occurs because biological rates are governed at the molecular level by the catalytic reaction rate, *k_cat_*. Based on transition state theory [3, 4]:

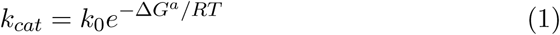

where Δ*G^a^* is the Gibbs free energy of activation required for the enzymatic reaction to occur (kcal mol*^−^*^1^), *R* is the universal gas constant (0.002 kcal mol*^−^*^1^), *T* is temperature measured in Kelvin, and *k*_0_ = *κk_B_T/h* where *κ* is a rate- and species-specific constant, *k_B_* is the Boltzmann constant, and *h* is Planck’s constant. Δ*G^a^* is comprised by an enthalpy term (Δ*H^a^*) and a temperature-dependent entropy term (Δ*S^a^*):

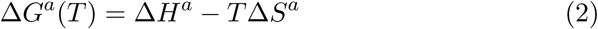

Warm temperature increases the entropy term (Δ*S^a^T* ), reducing Δ*G^a^*. Equations S1 and S2 thus describe an exponential increase in reaction rate with temperature.

### Strength of selection on catalytic rates and epistasis for fitness at cold temperatures

To form a hypothesis for how natural selection acts on catalytic rates at different temperatures, we incorporate ecological realism into predictions by assuming that the temperature-driven exponential increase in reaction rates will not have a 1:1 mapping with reproductive output, due to constraints on reproductive output set by other factors than temperature. At some point, suboptimal co-expression of correlated traits (evolutionary constraints), and/or limited nutrients or reproductive opportunities (ecological constraints) should make fitness pay-offs follow a pattern of diminishing returns with increases in catalytic rates [5–7] until temperature-dependent physiological rates are no longer the limiting factor on reproductive output (Fig. S1A). We describe these dynamics by assuming that reproductive rates are a logistic function of temperature, where the initial exponential increase in reproductive rate is explained by temperature-driven increases in catalytic rates and a release from thermodynamic constraints as temperatures warm, but where diminishing returns result from ecological constraints at higher temperatures:

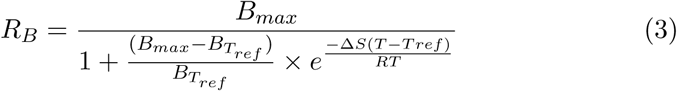

where *B_max_*is the maximum reproductive rate of the organism, and *B_Tref_* is reproductive output as a function of Δ*G^a^* at the reference temperature, *T_ref_* . We introduce a beneficial mutation that increases reproductive rate at the reference temperature (i.e., *B_Tref_* ) by 10% through effects on Δ*H^a^* (ΔΔ*H^a^* = −0.055 kcal/mol). We then estimate selection on this mutation across temperatures by comparing wildtype reproductive rate to that of the mutant (Fig. S1A, B). To estimate epistasis and the dependency of natural selection on genetic background, we performed these calculations on three genetic backgrounds with wildtype *B_Tref_* set to 40, 50 or 60 at *T_ref_* = 23°*C*, with *B_max_* = 100 in all cases. We further set Δ*S* to 0.50 kcal/mol K*^−^*^1^ based on values from the literature [8, 9].

This scenario generates strong positive selection on the introduced mutation across all three backgrounds at cold temperature (*s* = 0.1), while selection is weaker at intermediate benign temperature (*s* ≈ 0.005) and at hot temperature the mutation is effectively neutral (*s* ≈ 0) (Fig. S1B). This effect arises because cold temperatures act rate-limiting on reproductive output on all three backgrounds to an extent that far exceeds any ratelimiting ecological process. Once ecological constraints come into play at warmer temperatures, selection on the mutation weakens, but to similar extent across backgrounds, and little epistasis results.

### Strength of selection on protein stability and epistasis for fitness at hot temperatures

Organisms that experience temperatures that exceed their thermal optimum suffer from increased cellular stress and many physiological processes fail. This results in an exponential increase in molecular failure rates and cellular damage [1, 10]. One major component of cellular damage and decreased fitness at hot temperature is attributed to a reduction in the proportion of functional enzyme due to reversible inactivation via protein unfolding [11–15]. While this loss of protein stability is reversible, the reduction of properly folded protein leaves fewer molecules ready for work [14], and can cause excessive cellular toxicity as misfolded proteins clog up the cellular environment [16]. While several processes contribute to the observed exponential increase in death rates with temperature [1], here we use the example of protein folding stability to illustrate how thermodynamics predict temperature-dependent selection due to the rich empirical data allowing parameterization of the calculations. The proportion of folded enzyme ready to catalyse reactions follows a Boltzmann probability as a function of the Gibbs free energy of folding, Δ*G^f^* , which is thus a measure of protein stability [12]

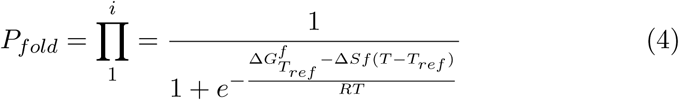

where stability (Δ*G^f^* ) is given for each of a set of *i* proteins that act in sequence and which effects are multiplicative [8, 14]. At a benign temperature of 25°C, most natural proteins occur in functional (properly folded) state, and mean Δ*G^f^* ≈ 7 kcal mol*^−^*^1^ [12, 17]. Warm temperatures increase entropy (Δ*S^f^* (*T* − *T_ref_* )) , reducing Δ*G^f^* , which leads to a rapid non-linear reduction in the fraction of functional enzyme (Fig. S1C). We introduce a beneficial mutation that increases folding stability at the reference temperature (*T_ref_*) and estimate selection on this mutation across temperatures by comparing the fraction of properly folded enzyme of the wildtype to that of the mutant (Fig. S1C). For the effect size of the mutation, we chose to increase folding stability by decreasing Δ*H^f^* by 1 kcal mol*^−^*^1^ (ΔΔ*H^f^* = −1.0 kcal/mol) based on empirical estimates [9, 17, 18]. We did this on three genetic backgrounds with the wildtype Δ*H^f^* in the enzyme receiving a mutation equal to 7, 9, or 11 kcal mol*^−^*^1^ at *T_ref_* = 23°C, which are typical stabilities observed across ectotherms (Dill et al. 2011). For other parameters we assumed Δ*S* = 0.50 kcal mol*^−^*^1^ K*^−^*^1^ and *i* = 100 proteins (e.g. [8, 17]).

In this scenario, the mutation acts as effectively neutral on all backgrounds at cold temperature (*s* ≈ 0). At intermediate temperature, there is selection on the background with the most unstable wildtype configuration (*s* ≈ 0.01) whereas the mutation is still effectively neutral on the other two backgrounds. At hot temperature the mutation is under strong selection on the backgrounds with low (*s >* 0.1) and intermediate (*s* ≈ 0.03) wildtype stability, whereas the mutation is still effectively neutral (*s* ≈ 0) on the background with most stable wildtype configuration (Fig. S1D), signifying substantial epistasis for fitness. This effect arises because proteins evolve marginal stability up until a point where selection is too weak to increase stability further [14, 19]. Around this zone of marginal stability, new mutations with conditional effects that are weak and effectively neutral at colder temperatures can accumulate, which affects subsequent selection on new genetic variants.

Protein folding can also be compromised by acute cold stress [1, 20]. However, such cold temperatures are typically not experienced during the active seasons of tropical and sub-tropical insects [21, 22], as the case for *C. maculatus* [23], and were therefore not modeled. Moreover, stressful temperatures affect organisms in a multitude of ways [1, 2, 5]. Thus, our particular example using protein folding is merely meant to illustrate a general principle and was chosen based on the simple fact that the modeled relationships are empirically well-supported in qualitative sense (see above). Other physiological phenomena that scale with temperature, such as metabolic expenditure [24] and the production of harmful reactive oxygen species [25], are likely to simultaneously contribute to increasing molecular failure rates at stressful temperature. If such processes follow similar patterns with temperature (i.e. marginally robust at the thermal optimum and governed by the same thermodynamics principles), similar predictions for the strength of selection and epistasis for fitness would result.

Yet, we have only illustrated predictions based on general, and very simplified, principles founded in thermodynamics. Our predictions assume multiplicative action between genes (or pathways) in effects on catalytic rates (Fig. S1A) and protein stability (Fig. S1C) that result in epistasis for fitness via non-linear (and temperature-dependent) mapping to fitness. We argue that these effects are general and may explain broad patterns in the average extent of epistasis for fitness across temperatures. However, at the more finegrained scale, molecular epistasis is the result of intricate and interdependent molecular processes which are likely to cause epistatic interactions among genes governing interacting pathways, which are likely omnipresent at any temperature.

**Figure S18:**
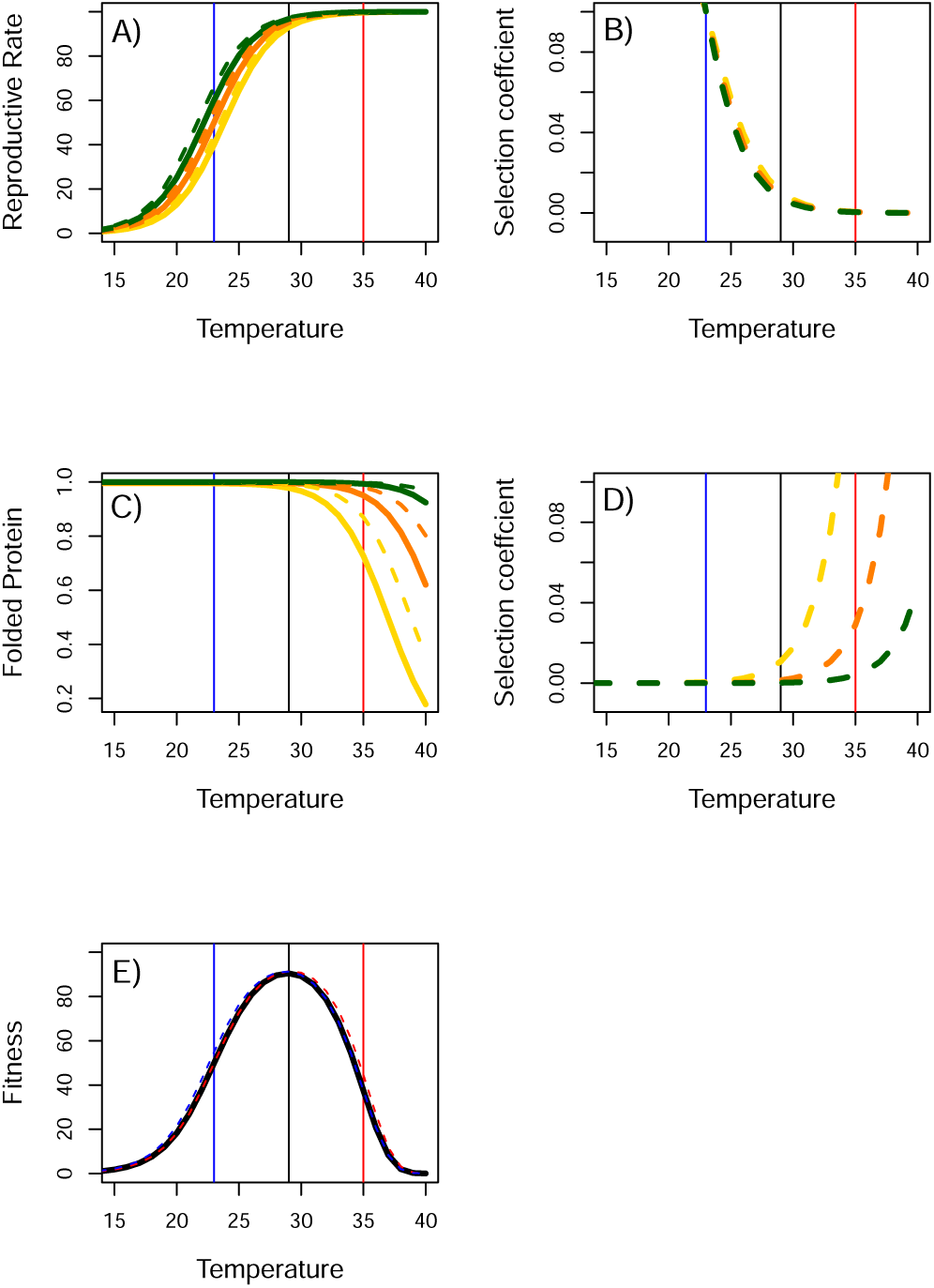
A hypothesis for temperature-dependent epistasis for fitness. A) Reproductive rate increases exponentially with temperature at the colder range due to relaxation of thermodynamics constraints on reaction rates, but starts to follow a pattern of diminishing returns at warmer temperatures due to ecological constraints. Illustrated for three genetic backgrounds (green, orange and yellow) with wildtypes (solid lines) and a mutant (broken lines) with a 10% increase in reproductive rate at 23°C. B) This results in strong selection on the mutation at cold temperature, but weak selection at hot temperature, and no epistasis. C) Warm temperatures increase molecular failure rates, for example via decreased protein stability. Illustrated for three genetic backgrounds with stable (green), intermediate (orange) and unstable (yellow) wildtype protein. A mutation increasing stability (broken lines) was introduced on each background. D) Selection on the mutation is weak at cold temperature for all backgrounds but can become under very strong selection at hot temperatures, depending on the wildtype protein stability, and strong epistasis for fitness results. E) Protein fitness as the product of reproductive rate and protein stability is depicted for the wildtype (black), and the mutants with increased reaction rate (blue) and increased protein stability (red), calculated for the yellow background.

## Notes

### Competing Interest Statement

The authors have declared no competing interest.

### Summary of Updates

Main file has been formatted to comply with journalistic guidelines (i.e., methods moved to the end). Some supplementary analyses have been added (e.g., AF-VapeR).

## Literature Cited

1. Lässig, M., Mustonen, V. & Walczak, A. M. Predicting evolution. Nature Ecology & Evolution 1, 0077 (2017).

2. Blount, Z. D., Lenski, R. E. & Losos, J. B. Contingency and determinism in evolution: Replaying life’s tape. Science 362, eaam5979 (2018).

3. Wortel, M. T., et al. Towards evolutionary predictions: Current promises and challenges. Evolutionary applications 16, 3–21 (2023).

4. Storz, J. F. Causes of molecular convergence and parallelism in protein evolution. Nature Reviews Genetics 17, 239 (2016).

5. Svensson, E. I. & Berger, D. The role of mutation bias in adaptive evolution. Trends in ecology & evolution 34, 422–434 (2019).

6. Orr, H. A. The probability of parallel evolution. Evolution 59, 216–220 (2005).

7. Chou, H.-H., Chiu, H.-C., Delaney, N. F., Segrè, D. & Marx, C. J. Diminishing returns epistasis among beneficial mutations decelerates adaptation. Science 332, 1190–1192 (2011).

8. Khan, A. I., Dinh, D. M., Schneider, D., Lenski, R. E. & Cooper, T. F. Negative epistasis between beneficial mutations in an evolving bacterial population. Science 332, 1193–1196 (2011).

9. Tenaillon, O., et al. The molecular diversity of adaptive convergence. Science 335, 457–461 (2012).

10. Kryazhimskiy, S., Rice, D. P., Jerison, E. R. & Desai, M. M. Global epistasis makes adaptation predictable despite sequence-level stochasticity. Science 344, 1519–1522 (2014).

11. Tenaillon, O., et al. Tempo and mode of genome evolution in a 50,000- generation experiment. Nature 536, 165–170 (2016).

12. Ono, J., Gerstein, A. C. & Otto, S. P. Widespread genetic incompatibilities between first-step mutations during parallel adaptation of Saccharomyces cerevisiae to a common environment. PLoS biology 15, e1002591 (2017).

13. Barghi, N., et al. Genetic redundancy fuels polygenic adaptation in Drosophila. PLoS biology 17, e3000128 (2019).

14. Whiting, J. R., et al. The genetic architecture of repeated local adaptation to climate in distantly related plants. Nature Ecology & Evolution, 1–15 (2024).

15. Hoffmann, A. A. & Sgro, C. M. Climate change and evolutionary adaptation. Nature 470, 479–485 (2011).

16. Hoffmann, A. A., Sgrò, C. M. & Kristensen, T. N. Revisiting adaptive potential, population size, and conservation. Trends in Ecology & Evolution 32, 506–517 (2017).

17. Capblancq, T., Fitzpatrick, M. C., Bay, R. A., Exposito-Alonso, M. & Keller, S. R. Genomic prediction of (mal) adaptation across current and future climatic landscapes. Annual Review of Ecology, Evolution, and Systematics 51, 245–269 (2020).

18. Waldvogel, A.-M., et al. Evolutionary genomics can improve prediction of species’ responses to climate change. Evolution Letters 4, 4–18 (2020).

19. Ghildiyal, K., et al. Genomic insights into the conservation of wild and animal diversity: a review. Gene, 147719 (2023).

20. Pearman, P. B., et al. Monitoring of species’ genetic diversity in Europe varies greatly and overlooks potential climate change impacts. Nature ecology & evolution 8, 267–281 (2024).

21. Hoffmann, A. A., Weeks, A. R. & Sgrò, C. M. Opportunities and challenges in assessing climate change vulnerability through genomics. Cell 184, 1420–1425 (2021).

22. Urban, M. C., et al. When and how can we predict adaptive responses to climate change? Evolution Letters 8, 172–187 (2024).

23. Conte, G. L., Arnegard, M. E., Peichel, C. L. & Schluter, D. The probability of genetic parallelism and convergence in natural populations. Proceedings of the Royal Society B: Biological Sciences 279, 5039–5047 (2012).

24. Burny, C., Nolte, V., Dolezal, M. & Schlötterer, C. Highly parallel genomic selection response in replicated Drosophila melanogaster populations with reduced genetic variation. Genome Biology and Evolution 13, evab239 (2021).

25. Schlötterer, C. How predictable is adaptation from standing genetic variation? Experimental evolution in Drosophila highlights the central role of redundancy and linkage disequilibrium. Philosophical Transactions of the Royal Society B 378, 20220046 (2023).

26. Weinreich, D. M., Delaney, N. F., DePristo, M. A. & Hartl, D. L. Darwinian evolution can follow only very few mutational paths to fitter proteins. science 312, 111–114 (2006).

27. De Visser, J. A. G., Cooper, T. F. & Elena, S. F. The causes of epistasis. Proceedings of the Royal Society B: Biological Sciences 278, 3617–3624 (2011).

28. Bohutínská, M., et al. Genomic basis of parallel adaptation varies with divergence in Arabidopsis and its relatives. Proceedings of the National Academy of Sciences 118, e2022713118 (2021).

29. Agrawal, A. F. & Whitlock, M. C. Environmental duress and epistasis: how does stress affect the strength of selection on new mutations? Trends in ecology & evolution 25, 450–458 (2010).

30. Caruso, C. M., et al. What are the environmental determinants of phenotypic selection? A meta-analysis of experimental studies. The American Naturalist 190, 363–376 (2017).

31. Hutter, S., Saminadin-Peter, S. S., Stephan, W. & Parsch, J. Gene expression variation in African and European populations of Drosophila melanogaster. Genome biology 9, 1–15 (2008).

32. Mallard, F., Nolte, V., Tobler, R., Kapun, M. & Schlötterer, C. A simple genetic basis of adaptation to a novel thermal environment results in complex metabolic rewiring in Drosophila. Genome biology 19, 1–15 (2018).

33. Somero, G. N., Lockwood, B. L. & Tomanek, L. Biochemical adaptation: Response to Environmental Challenges from Life’s Origins to the Anthropocene (Oxford university press, 2017).

34. Angilletta, M. J. Thermal adaptation: a theoretical and empirical synthesis (Oxford University Press, 2009).

35. Huey, R. B. & Kingsolver, J. G. Evolution of thermal sensitivity of ectotherm performance. Trends in ecology & evolution 4, 131–135 (1989).

36. Drake, J. W. Avoiding dangerous missense: thermophiles display especially low mutation rates. PLoS genetics 5, e1000520 (2009).

37. Agozzino, L. & Dill, K. A. Protein evolution speed depends on its stability and abundance and on chaperone concentrations. Proceedings of the National Academy of Sciences 115, 9092–9097 (2018).

38. Berger, D., Stångberg, J., Baur, J. & Walters, R. J. Elevated temperature increases genome-wide selection on de novo mutations. Proceedings of the Royal Society B 288, 20203094 (2021).

39. Burc, E., et al. Life-history adaptation under climate warming magnifies the agricultural footprint of a cosmopolitan insect pest. bioRxiv, 2024–03 (2024).

40. Krill, J. L. & Dawson-Scully, K. cGMP-dependent protein kinase inhibition extends the upper temperature limit of stimulus-evoked calcium responses in motoneuronal boutons of Drosophila melanogaster larvae. Plos one 11, e0164114 (2016).

41. Schmidt, P. S., et al. An amino acid polymorphism in the couch potato gene forms the basis for climatic adaptation in Drosophila melanogaster. Proceedings of the National Academy of Sciences 105, 16207–16211 (2008).

42. Zhang, Q. & Denlinger, D. L. Elevated couch potato transcripts associated with adult diapause in the mosquito Culex pipiens. Journal of insect physiology 57, 620–627 (2011).

43. Sakamoto, F., et al. Detection of evolutionary conserved and accelerated genomic regions related to adaptation to thermal niches in Anolis lizards. Ecology and Evolution 14, e11117 (2024).

44. Gerken, A. R., Eller, O. C., Hahn, D. A. & Morgan, T. J. Constraints, independence, and evolution of thermal plasticity: probing genetic architecture of long-and short-term thermal acclimation. Proceedings of the National Academy of Sciences 112, 4399–4404 (2015).

45. Fang, Y., Gong, H., Yang, R., Lai, K. W. & Quan, M. An active biomechanical model of cell adhesion actuated by intracellular tensioningtaxis. Biophysical journal 118, 2656–2669 (2020).

46. Das, S. G. & Krug, J. Unpredictable repeatability in molecular evolution. Proceedings of the National Academy of Sciences 119, e2209373119 (2022).

47. Diaz-Colunga, J., Sanchez, A. & Ogbunugafor, C. B. Environmental modulation of global epistasis in a drug resistance fitness landscape. Nature Communications 14, 8055 (2023).

48. Echave, J. & Wilke, C. O. Biophysical models of protein evolution: understanding the patterns of evolutionary sequence divergence. Annual review of biophysics 46, 85–103 (2017).

49. Chen, P. & Shakhnovich, E. I. Thermal adaptation of viruses and bacteria. Biophysical journal 98, 1109–1118 (2010).

50. Bloom, J. D., Labthavikul, S. T., Otey, C. R. & Arnold, F. H. Protein stability promotes evolvability. Proceedings of the National Academy of Sciences 103, 5869–5874 (2006).

51. Goldstein, R. A. The evolution and evolutionary consequences of marginal thermostability in proteins. Proteins: Structure, Function, and Bioinformatics 79, 1396–1407 (2011).

52. Gain, C., et al. A quantitative theory for genomic offset statistics. Molecular Biology and Evolution 40, k(2023).

53. Lotterhos, K. E. Interpretation issues with “genomic vulnerability” arise from conceptual issues in local adaptation and maladaptation. Evolution Letters 8, 331–339 (2024).

54. Parmesan, C. Ecological and evolutionary responses to recent climate change. Annu. Rev. Ecol. Evol. Syst. 37, 637–669 (2006).

55. Urban, M. C. Accelerating extinction risk from climate change. Science 348, 571–573 (2015).

56. Radchuk, V., et al. Adaptive responses of animals to climate change are most likely insufficient. Nature communications 10, 3109 (2019).

57. Rellstab, C., Dauphin, B. & Exposito-Alonso, M. Prospects and limitations of genomic offset in conservation management. Evolutionary Applications 14, 1202–1212 (2021).

58. Barton, N. H. The “new synthesis”. Proceedings of the National Academy of Sciences 119, e2122147119 (2022).

59. Echave, J., Spielman, S. J. & Wilke, C. O. Causes of evolutionary rate variation among protein sites. Nature Reviews Genetics 17, 109–121 (2016).

60. Bloom, J. D., et al. Thermodynamic prediction of protein neutrality. Proceedings of the National Academy of Sciences 102, 606–611 (2005).

61. Tobler, R., et al. Massive habitat-specific genomic response in D. melanogaster populations during experimental evolution in hot and cold environments. Molecular biology and evolution 31, 364–375 (2014).

62. Otte, K. A., Nolte, V., Mallard, F. & Schlötterer, C. The genetic architecture of temperature adaptation is shaped by population ancestry and not by selection regime. Genome Biology 22, 1–25 (2021).

63. Hayward, L. K. & Sella, G. Polygenic adaptation after a sudden change in environment. Elife 11, e66697 (2022).

64. Gallegos, C., Hodgins, K. A. & Monro, K. Climate adaptation and vulnerability of foundation species in a global change hotspot. Molecular Ecology 32, 1990–2004 (2023).

65. Van Boheemen, L. A. & Hodgins, K. A. Rapid repeatable phenotypic and genomic adaptation following multiple introductions. Molecular Ecology 29, 4102–4117 (2020).

66. Renaut, S., Owens, G. L. & Rieseberg, L. H. Shared selective pressure and local genomic landscape lead to repeatable patterns of genomic divergence in sunflowers. Molecular Ecology 23, 311–324 (2014).

67. Chaturvedi, S., et al. Climatic similarity and genomic background shape the extent of parallel adaptation in Timema stick insects. Nature ecology & evolution 6, 1952–1964 (2022).

68. Feder, M. E., Bennett, A. F. & Huey, R. B. Evolutionary physiology. Annual review of ecology and systematics 31, 315–341 (2000).

69. Weinreich, D. M., Watson, R. A. & Chao, L. Perspective: sign epistasis and genetic costraint on evolutionary trajectories. Evolution 59, 1165–1174 (2005).

70. Bank, C. Epistasis and adaptation on fitness landscapes. Annual Review of Ecology, Evolution, and Systematics 53, 457–479 (2022).

71. Li, C. & Zhang, J. Multi-environment fitness landscapes of a tRNA gene. Nature ecology & evolution 2, 1025–1032 (2018).

72. Johnson, M. S., Reddy, G. & Desai, M. M. Epistasis and evolution: recent advances and an outlook for prediction. BMC biology 21, 120 (2023).

73. Fitzpatrick, M. C., Chhatre, V. E., Soolanayakanahally, R. Y. & Keller, S. R. Experimental support for genomic prediction of climate maladaptation using the machine learning approach Gradient Forests. Molecular Ecology Resources 21, 2749–2765 (2021).

74. De Lisle, S. P. & Bolnick, D. I. A multivariate view of parallel evolution. Evolution 74, 1466–1481 (2020).

75. Bailey, S. F., Rodrigue, N. & Kassen, R. The effect of selection environment on the probability of parallel evolution. Molecular biology and evolution 32, 1436–1448 (2015).

76. Guedes, R. N., Guedes, N. M. & Smith, R. H. Larval competition within seeds: from the behaviour process to the ecological outcome in the seed beetle Callosobruchus maculatus. Austral Ecology 32, 697–707 (2007).

77. Santos, M. A., et al. Experimental Evolution in a Warming World: The Omics Era. Molecular Biology and Evolution 41, msae148 (2024).

78. Czech, L., et al. Monitoring rapid evolution of plant populations at scale with Pool-Sequencing. BioRxiv, 2022–02 (2022).

79. Tuda, M., Kagoshima, K., Toquenaga, Y. & Arnqvist, G. Global genetic differentiation in a cosmopolitan pest of stored beans: effects of geography, host-plant usage and anthropogenic factors. PLoS One 9, e106268 (2014).

80. Immonen, E., Berger, D., Sayadi, A., Liljestrand-Rönn, J. & Arnqvist, G. An experimental test of temperature-dependent selection on mitochondrial haplotypes in Callosobruchus maculatus seed beetles. Ecology and evolution 10, 11387–11398 (2020).

81. Dowling, D. K., Abiega, K. C. & Arnqvist, G. Temperature-specific outcomes of cytoplasmic-nuclear interactions on egg-to-adult development time in seed beetles. Evolution 61, 194–201 (2007).

82. Dowling, D. K., Friberg, U. & Arnqvist, G. A comparison of nuclear and cytoplasmic genetic effects on sperm competitiveness and female remating in a seed beetle. Journal of Evolutionary Biology 20, 2113–2125 (2007).

83. Berger, D. & Liljestrand-Rönn, J. Environmental complexity mitigates the demographic impact of sexual selection. Ecology Letters 27, e14355 (2024).

84. Bolnick, D. I., Barrett, R. D., Oke, K. B., Rennison, D. J. & Stuart, Y. E. (Non) parallel evolution. Annual Review of Ecology, Evolution, and Systematics 49, 303–330 (2018).

85. Hansen, T. F. & Houle, D. Measuring and comparing evolvability and constraint in multivariate characters. Journal of evolutionary biology 21, 1201–1219 (2008).

86. Berger, D., Postma, E., Blanckenhorn, W. U. & Walters, R. J. Quantitative genetic divergence and standing genetic (co) variance in thermal reaction norms along latitude. Evolution 67, 2385–2399 (2013).

87. Li, H. & Durbin, R. Fast and accurate short read alignment with Burrows–Wheeler transform. bioinformatics 25, 1754–1760 (2009).

88. McKenna, A., et al. The Genome Analysis Toolkit: a MapReduce framework for analyzing next-generation DNA sequencing data. Genome research 20, 1297–1303 (2010).

89. Pockrandt, C., Alzamel, M., Iliopoulos, C. S. & Reinert, K. GenMap: ultra-fast computation of genome mappability. Bioinformatics 36, 3687–3692 (2020).

90. Jónás, Á., Taus, T., Kosiol, C., Schlötterer, C. & Futschik, A. Estimating the effective population size from temporal allele frequency changes in experimental evolution. Genetics 204, 723–735 (2016).

91. Ewens, W. J. Mathematical Population Genetics I: Theoretical Introduction (Springer Science & Business Media, 2012).

92. Gompert, Z. & Messina, F. J. Genomic evidence that resource-based trade-offs limit host-range expansion in a seed beetle. Evolution 70, 1249–1264 (2016).

93. Iranmehr, A., Akbari, A., Schlötterer, C. & Bafna, V. CLEAR: Composition of likelihoods for evolve and resequence experiments. Genetics 206, 1011–1023 (2017).

94. Kofler, R., et al. PoPoolation: a toolbox for population genetic analysis of next generation sequencing data from pooled individuals. PloS one 6, e15925 (2011).

95. Whiting, J. R., Paris, J. R., van der Zee, M. J. & Fraser, B. A. AF- vapeR: a multivariate genome scan for detecting parallel evolution using allele frequency change vectors. Methods in Ecology and Evolution 13, 2167–2180 (2022).

## Supplementary Literature Cited

1. Somero, G. N., Lockwood, B. L. & Tomanek, L. Biochemical adaptation: Response to Environmental Challenges from Life’s Origins to the Anthropocene (Oxford university press, 2017).

2. Angilletta, M. J. Thermal adaptation: a theoretical and empirical synthesis (Oxford University Press, 2009).

3. Evans, M. G. & Polanyi, M. Some applications of the transition state method to the calculation of reaction velocities, especially in solution. Transactions of the Faraday Society 31, 875–894 (1935).

4. Eyring, H. The activated complex in chemical reactions. The Journal of Chemical Physics 3, 107–115 (1935).

5. Clarke, A. Is there a universal temperature dependence of metabolism? Functional Ecology 18, 252–256 (2004).

6. Berger, D., Walters, R. & Gotthard, K. What limits insect fecundity? Body size-and temperature-dependent egg maturation and oviposition in a butterfly. Functional Ecology 22, 523–529 (2008).

7. Dell, A. I., Pawar, S. & Savage, V. M. Systematic variation in the temperature dependence of physiological and ecological traits. Proceedings of the National Academy of Sciences 108, 10591–10596 (2011).

8. Chen, P. & Shakhnovich, E. I. Thermal adaptation of viruses and bacteria. Biophysical journal 98, 1109–1118 (2010).

9. Dill, K. A., Ghosh, K. & Schmit, J. D. Physical limits of cells and proteomes. Proceedings of the National Academy of Sciences 108, 17876–17882 (2011).

10. Jørgensen, L. B., Ørsted, M., Malte, H., Wang, T. & Overgaard, J. Extreme escalation of heat failure rates in ectotherms with global warming. Nature 611, 93–98 (2022).

11. Bloom, J. D., et al. Thermodynamic prediction of protein neutrality. Proceedings of the National Academy of Sciences 102, 606–611 (2005).

12. DePristo, M. A., Weinreich, D. M. & Hartl, D. L. Missense meanderings in sequence space: a biophysical view of protein evolution. Nature Reviews Genetics 6, 678–687 (2005).

13. Bershtein, S., Serohijos, A. W. & Shakhnovich, E. I. Bridging the physical scales in evolutionary biology: from protein sequence space to fitness of organisms and populations. Current opinion in structural biology 42, 31–40 (2017).

14. Echave, J. & Wilke, C. O. Biophysical models of protein evolution: understanding the patterns of evolutionary sequence divergence. Annual review of biophysics 46, 85–103 (2017).

15. Agozzino, L. & Dill, K. A. Protein evolution speed depends on its stability and abundance and on chaperone concentrations. Proceedings of the National Academy of Sciences 115, 9092–9097 (2018).

16. Allan Drummond, D. & Wilke, C. O. The evolutionary consequences of erroneous protein synthesis. Nature Reviews Genetics 10, 715–724 (2009).

17. Chen, P. & Shakhnovich, E. I. Lethal mutagenesis in viruses and bacteria. Genetics 183, 639–650 (2009).

18. Zeldovich, K. B., Chen, P. & Shakhnovich, E. I. Protein stability imposes limits on organism complexity and speed of molecular evolution. Proceedings of the National Academy of Sciences 104, 16152–16157 (2007).

19. Goldstein, R. A. The evolution and evolutionary consequences of marginal thermostability in proteins. *Proteins: Structure*, Function, and Bioinformatics 79, 1396–1407 (2011).

20. Feder, M. E., Bennett, A. F. & Huey, R. B. Evolutionary physiology. Annual review of ecology and systematics 31, 315–341 (2000).

21. Deutsch, C. A., et al. Impacts of climate warming on terrestrial ectotherms across latitude. Proceedings of the National Academy of Sciences 105, 6668–6672 (2008).

22. Johansson, F., Orizaola, G. & Nilsson-Örtman, V. Temperate insects with narrow seasonal activity periods can be as vulnerable to climate change as tropical insect species. Scientific Reports 10, 8822 (2020).

23. Baur, J., Zwoinska, M., Koppik, M., Snook, R. R. & Berger, D. Heat stress reveals a fertility debt owing to postcopulatory sexual selection. Evolution Letters 8, 101–113 (2024).

24. Gillooly, J. F., Brown, J. H., West, G. B., Savage, V. M. & Charnov, E. L. Effects of size and temperature on metabolic rate. science 293, 2248–2251 (2001).

25. Dowling, D. K. & Simmons, L. W. Reactive oxygen species as universal constraints in life-history evolution. Proceedings of the Royal Society B: Biological Sciences 276, 1737–1745 (2009).

